# Opposing functions for retromer and Rab11 in extracellular vesicle cargo traffic at presynaptic terminals

**DOI:** 10.1101/645713

**Authors:** Rylie B. Walsh, Agata N. Becalska, Matthew J. Zunitch, Tania Lemos, Erica C. Dresselhaus, So Min Lee, ShiYu Wang, Berith Isaac, Anna Yeh, Kate Koles, Avital A. Rodal

**Affiliations:** Department of Biology, Brandeis University, Waltham, MA 02453

**Keywords:** Amyloid Precursor Protein, Synaptotagmin-4, retromer, Vps35, Rab11, *Drosophila*, extracellular vesicle, synapse, endosome, exosome

## Abstract

Neuronal extracellular vesicles (EVs) play important roles in intercellular communication and pathogenic protein propagation in neurological disease. However, it remains unclear how cargoes are selectively packaged into neuronal EVs. Here, we show that loss of the endosomal retromer complex leads to accumulation of EV cargoes Amyloid Precursor Protein (APP) and Synaptotagmin-4 (Syt4) at *Drosophila* motor neuron presynaptic terminals, resulting in increased release of these cargoes in EVs. By systematically exploring known retromer-dependent trafficking mechanisms, we show that EV regulation is separable from several previously identified roles of neuronal retromer, and depends on the ESCPE-1 complex. Conversely, loss of the recycling endosome regulator *rab11* leads to reduced EV cargo levels, and suppresses cargo accumulation in retromer mutants. Thus, EV traffic reflects a balance between Rab11-mediated loading and retromer-dependent removal from EV precursor compartments. Our data shed light on previous studies implicating Rab11 and retromer in competing pathways in Alzheimer’s Disease, and suggest that misregulated EV traffic may be an underlying defect.

## INTRODUCTION

Transmembrane proteins are routed from the plasma membrane through an intersecting set of endolysosomal trafficking pathways that determine whether proteins are degraded or recycled (Naslavsky and Caplan, 2018). Effective sorting between these endosomal pathways is particularly critical for the survival and function of neurons, and aberrant endosomal trafficking is linked to many neurodegenerative diseases, including Alzheimer’s Disease (AD) (Small et al., 2017; Winckler et al., 2018). One possible fate for endosomal cargo is incorporation into small secreted vesicles called extracellular vesicles (EVs). EVs include exosomes (derived when endosomal multivesicular bodies (MVBs) fuse with the plasma membrane, releasing their intralumenal vesicles (ILVs) into the extracellular space) and microvesicles (derived from direct budding from the plasma membrane) (van Niel et al., 2018). EVs can contain signaling proteins, lipids, and nucleic acids, and serve as an important means of communication between different cell types in the nervous system (Holm et al., 2018). Neuronal EVs have also recently received attention as a potential mechanism for both intercellular propagation and clearance of pathogenic protein species, including the Amyloid Precursor Protein (APP), a central player in AD (Arbo et al., 2020; Song et al., 2020). APP is a type I transmembrane protein that traffics through the endosomal system, where it can be proteolytically processed into fragments including toxic Amyloid-β (Aβ a major constituent of the hallmark amyloid plaques of AD (Tan and Gleeson, 2019). APP C-terminal fragments and Aβ have been found in pathogenic EVs (Laulagnier et al., 2017; Perez-Gonzalez et al., 2012; Rajendran et al., 2006; Sardar Sinha et al., 2018; Sharples et al., 2008), but the mechanisms by which APP is sorted into EVs are not well understood, and are challenging to study *in vivo* at synapses, which may be important sites of Aβ release (Dolev et al., 2013; Lazarov et al., 2002; Lundgren et al., 2014; Schedin-Weiss et al., 2016; Yu et al., 2018).

Retromer, a heterotrimeric complex composed of Vps35, Vps26, and Vps29, is a central component of the endosomal trafficking machinery, and mediates cargo sorting into membrane tubules and buds (Chen et al., 2019a; McNally and Cullen, 2018; Wang et al., 2018). Retromer serves a wide variety of cellular functions, including recycling numerous cargoes from endosomes to the Golgi, plasma membrane, or secretory granules (Wang et al., 2018; Neuman et al., 2020), controlling endosome maturation (Jimenez-Orgaz et al., 2017; Kvainickas et al., 2019; Seaman et al., 2018; Ye et al., 2020), retrieving lysosomal/vacuolar hydrolase receptors to protect them from degradation (Arighi et al., 2004; Cui et al., 2019; Seaman et al., 1998), facilitating autophagy (Jimenez-Orgaz et al., 2017), and influencing mitochondria fusion and function (Tang et al., 2015). The role of retromer in endosome-derived EV traffic is less clear. While depletion of retromer leads to increased APP in EVs released from HEK293T cells (Sullivan et al., 2011), others have found decreased EV levels of Wnt3A and its carrier Evi/Wntless upon loss of retromer from this same cell type (Gross et al., 2012), and little is known about the role of retromer in EV traffic in neurons.

Neuronal retromer localizes to cell bodies, dendrites, and axons, and is required for proper synaptic morphology, synaptic transmission, synaptic vesicle number, and AMPA receptor traffic (Bhalla et al., 2012; Choy et al., 2014; Inoshita et al., 2017; Korolchuk et al., 2007; Munsie et al., 2015; Temkin et al., 2017; Tian et al., 2015; Vazquez-Sanchez et al., 2018; Wu et al., 2017). Retromer levels are reduced in the entorhinal cortex of AD patients (Small et al., 2005), and it has therefore been proposed as a therapeutic target (Berman et al., 2015; Mecozzi et al., 2014). Indeed, in cellular and animal models of AD, loss of retromer leads to increased Aβ, exacerbates synaptic and cognitive defects, and causes mislocalization of APP as well as its amyloidogenic protease BACE1 (Bhalla et al., 2012; Choy et al., 2012; Eggert et al., 2018; Li et al., 2019; Muhammad et al., 2008; Neuman et al., 2020; Sullivan et al., 2011; Tan and Gleeson, 2019; Wen et al., 2011). However, despite the importance of retromer in AD and endosome biology, and of endosome-derived EVs in neurological function and disease, we do not yet understand if and how neuronal retromer influences the traffic of APP and/or other cargoes into EVs. This will be particularly important for retromer-directed therapies, which should target specific pathological functions of retromer rather than all of its functions.

Here, we test the hypothesis that retromer is involved in sorting neuronal EV cargo including APP, taking advantage of the highly accessible presynaptic terminals at the *Drosophila* larval neuromuscular junction (NMJ), which provide a powerful system for studying EVs in their *in vivo* context in the animal (Koles et al., 2012; Korkut et al., 2009; Korkut et al., 2013).

## RESULTS

### Neuronally-expressed APP traffics into EVs

To examine sub-cellular trafficking of APP at *Drosophila* synapses, we generated transgenic flies expressing human APP^695^ cDNA tagged at its C terminus with EGFP, under the control of the binary GAL4-UAS system (**Fig. 1A**). Neuronally-driven APP-GFP localized to motile particles in segmental axons of third instar larvae, as previously reported (Gunawardena and Goldstein, 2001; Ramaker et al., 2016). Surprisingly, in addition to its expected presynaptic localization at the NMJ, we found a large fraction of neuronally-expressed APP-EGFP in stationary extra-neuronal postsynaptic puncta (**Fig 1B**). Since the EGFP tag is located on the intracellular C-terminus of APP, these extra-neuronal puncta are likely encapsulated by neuronally-derived membrane, as opposed to representing the shed extracellular domain of APP. In support of this hypothesis, postsynaptic APP-EGFP puncta colocalized with α-HRP (which recognizes a number of neuronal glycoproteins (Snow et al., 1987)), and with the EV markers Tsp42Ej (the *Drosophila* homolog of mammalian tetraspanin CD63) and Evi/Wntless (Korkut et al., 2009) (**Fig. 1C, D**). By contrast, a cytosolic presynaptic protein, Complexin (Cpx), was excluded from postsynaptic APP-EGFP puncta, indicating that they arise from specific presynaptic sorting events rather than from non-specific shedding of presynaptic membrane and cytoplasm (**Fig. 1B**). An antibody against amino acids 17-24 of the APP Aβ fragment also co-localized with APP-EGFP, indicating that these puncta contain membrane-spanning APP sequences (**Fig. 1D**). By contrast, APP-EGFP expressed with a muscle GAL4 driver did not accumulate in α-HRP-positive postsynaptic puncta, and instead was diffusely located within the muscle subsynaptic reticulum (SSR), indicating that postsynaptic puncta of presynaptically-driven APP-EGFP did not arise from its leaky expression in muscles (**Fig. S1A**). Thus, a large proportion of presynaptic APP is trafficked into postsynaptic EV-like structures that are associated with the neuronal membrane.

**Fig. 1.**
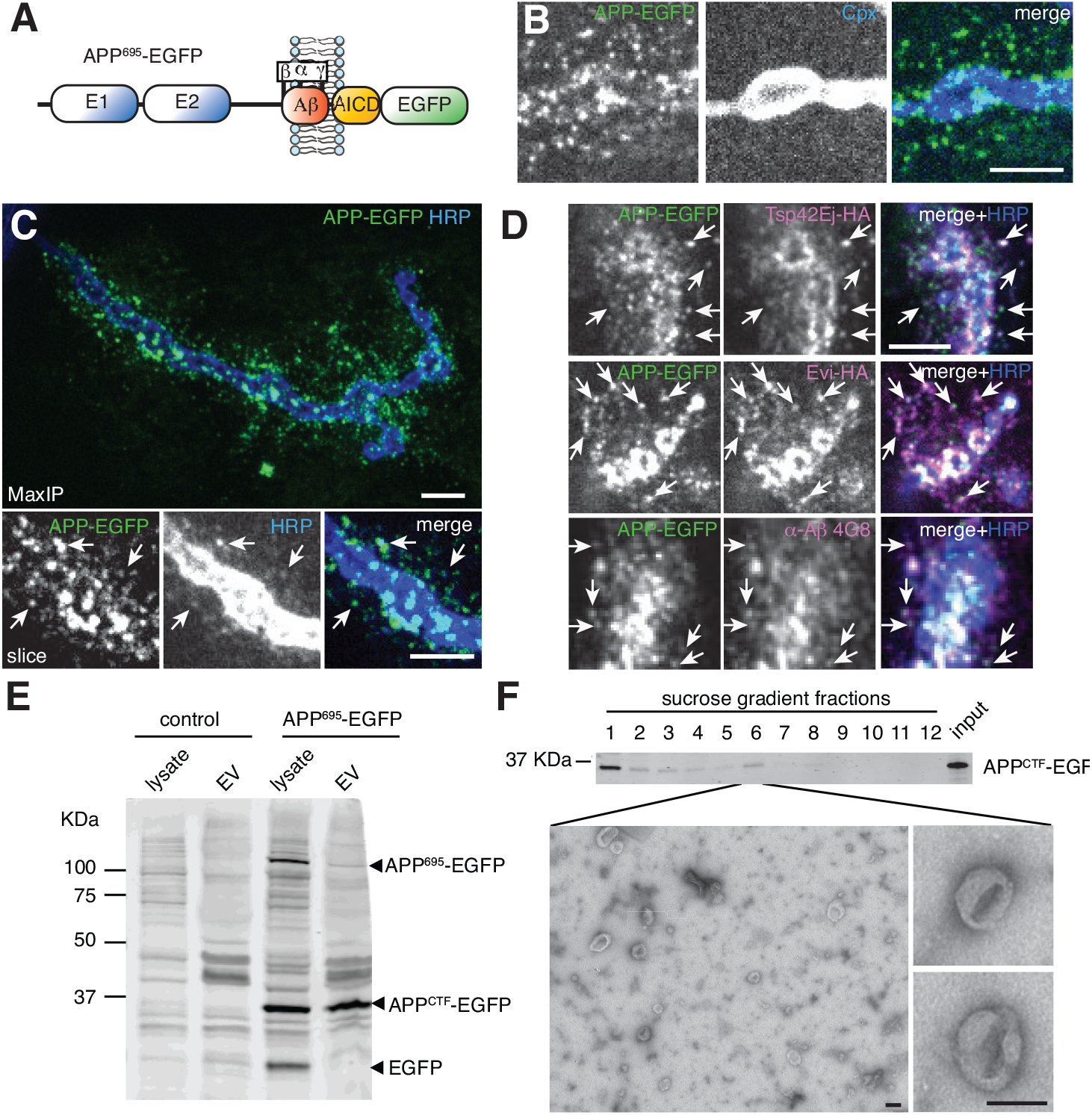
APP localizes to neuronally-derived EVs. (**A**) Schematic of APP-EGFP construct. (**B-D**) Spinning disk confocal images from 3^rd^ instar larval muscle 4 NMJs expressing neuronally-driven (GAL4^C155^) UAS-APP-EGFP and labeled with the indicated antibodies. White arrows highlight examples of co-localization. Scale bars are 5 µm. (**B**) Maximum intensity projection (MaxIP) of motor-neuron-derived APP-EGFP localization presynaptically and to extraneuronal puncta that exclude the presynaptic cytoplasmic protein Complexin (Cpx). (**C**) (MaxIP) Motor-neuron-derived post-synaptic APP-EGFP co-localization with the neuronal membrane marker α-HRP (**D**) (Single confocal slices) Motor-neuron-derived APP-EGFP co-localization with neuronally-expressed EV cargoes Tsp42Ej and Evi, as well as a transmembrane epitope in APP. (**E,F**) α-GFP immunoblots of fractions from S2 cells stably expressing APP-EGFP. (**F**) Negative-stain EM of an APP CTF-containing 1.14 mg/ml sucrose gradient fraction. Scale bar is 100 nm. **See also Figure S1.**

We next confirmed that APP could sort to *bona fide* EVs in other cell types. We generated a stable *Drosophila* S2 cell line expressing APP-EGFP, fractionated cell culture supernatants, and immunoblotted the EV fraction for APP-EGFP. EVs have been demonstrated to pellet from cell culture supernatant at high speed (100,000 x *g*), fractionate on equilibrium gradients at 1.1-1.2 mg/ml sucrose, and exhibit a typical cup-shaped morphology by negative stain electron microscopy (Thery et al., 2018). We found that the purified EV fraction from S2 cell supernatants met these criteria (**Fig 1E, F**). Notably, the ratio of an EGFP-tagged C-terminal fragment of APP (CTF) compared to full-length APP was greatly enhanced in the S2 cell EV fraction relative to total cell extracts, while EGFP alone did not fractionate with EVs. Indeed, S2 cells have been previously shown to express proteases capable of processing APP family members (Luo et al., 1990), and similar CTFs of endogenous APP are enriched in EVs derived from cortical neurons (Laulagnier et al., 2017). CTFs are generated by endosomally localized proteases (Tan and Gleeson, 2019), suggesting that APP EVs are endosomally derived exosomes. Taken together, these results indicate that APP-EGFP, and particularly a CTF of this protein, are trafficked into EVs by multiple established criteria (Thery et al., 2018).

We next tested whether EV localization at synapses is conserved for Appl, the *Drosophila* homolog of APP, which is expressed in neurons and acts as a signaling factor in neuronal development and maintenance (Cassar and Kretzschmar, 2016). In wild-type animals, Appl α-C-term antibodies (Swanson et al., 2005) labeled postsynaptic puncta that co-localized with α-HRP-positive neuronally-derived membrane. These puncta were more prevalent in animals presynaptically overexpressing UAS-Appl. By contrast, non-specific postsynaptic puncta in *appld* null mutants did not co-localize with α-HRP (**Fig. S1C**). This suggests that like APP-EGFP, Appl containing its intracellular C terminus is trafficked into EVs derived from neuronal membranes. Further supporting that postsynaptic puncta represent Appl EV localization rather than ectodomain shedding, Appl α-C-term antibodies labeled postsynaptic puncta for neuronally expressed cleavage-resistant Appl (Appl^SD^ (Torroja et al., 1999)), but not Appl^SD^ lacking the C-terminal intracellular antibody epitope (**Fig. S1D**). These results indicate that like human APP-GFP, *Drosophila* Appl is trafficked into EVs, and that EVs are a normal trajectory for neuronal APP *in vivo*.

### Neuronal retromer restricts accumulation of EV cargoes

At the *Drosophila* NMJ, retromer regulates synaptic vesicle size and recycling, as well as synaptic growth (Inoshita et al.; Korolchuk et al., 2007; Ye et al., 2020). Retromer could potentially also be involved in EV-mediated release of APP from neurons, as had been suggested previously in HEK293T cells (Sullivan et al., 2011), but this hypothesis has not yet been tested at synapses *in vivo*. To address this question, we measured APP-EGFP intensity in the presynaptic neuron and the surrounding postsynaptic region in *Vps35* mutants. We found that the mean intensity of APP-EGFP was strongly increased at *Vps35* mutant NMJs, in both presynaptic and postsynaptic puncta, and in the overall presynaptic volume (**Fig. 2A-C**). We found a similar increase in APP-EGFP levels in cell bodies and axons (**Fig. 2D-E**). Since synapses are a site of EV release in these motor neurons (Korkut et al., 2009; Korkut et al., 2013), we focused our subsequent studies on the mechanisms by which *Vps35* mutants lead to cargo accumulation and release from this site. Using structured illumination microscopy (SIM), we resolved individual presynaptic puncta and found that increased APP-EGFP levels occurred as larger and more intense puncta, rather than as a greater density of puncta (**Fig S2A).**Importantly, the overall increase in APP levels was insufficient to explain the greater postsynaptic APP EV abundance: At both normal and *Vps35* mutant NMJs, the fraction of postsynaptic APP did not correlate strongly with total (presynaptic * postsynaptic) APP levels (**Fig S2B**). Further, the ratio of postsynaptic to total APP was significantly increased in *Vps35* mutants (**Fig S2C**). These data suggest that the EV phenotype of Vps35 mutants is mediated by a specific EV-regulatory pathway rather than simply by increased overall APP levels.

**Figure 2.**
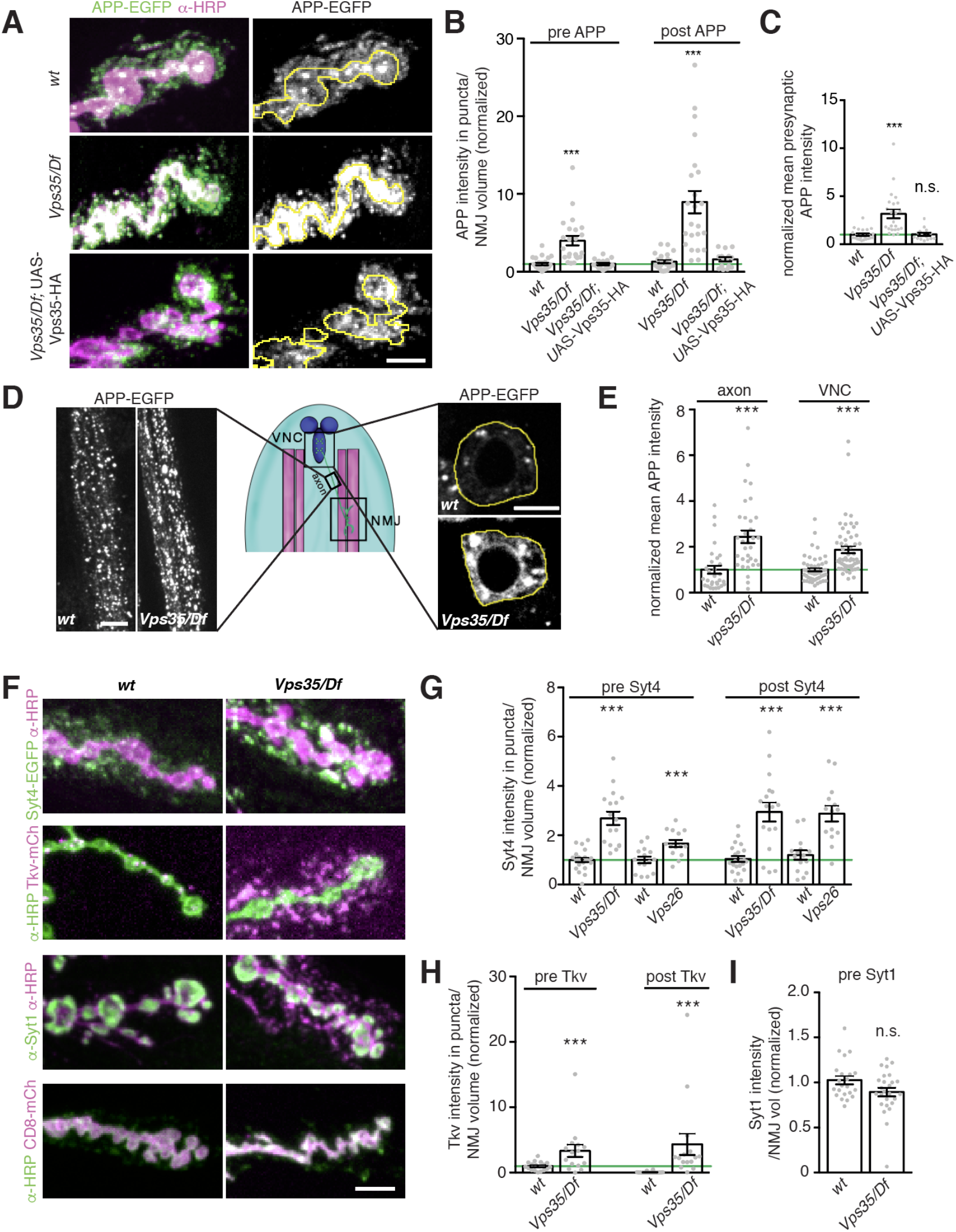
Neuronal retromer restricts accumulation of cargoes presynaptically and in EVs. **(A-C)** Loss of *Vps35* causes accumulation of presynaptic and postsynaptic APP-EGFP, both of which are rescued by presynaptic expression of Vps35-HA. **(A)**MaxIPs of NMJs expressing neuronal driver GAL4^C155^ and UAS-APP-EGFP in the indicated genotypes. (**B**) Quantification of presynaptic and postsynaptic (within 3 µm of presynaptic membrane) intensity of APP-EGFP in thresholded puncta. (**C**) Overall (unthresholded) APP-EGFP intensity in the presynaptic volume is increased in *Vps35* mutants and rescued by presynaptic expression of Vps35-HA. (**D,E**) APP-EGFP intensity increases in the axons and cell bodies of *Vps35* mutant motor neurons. (**D**) Single confocal slices of APP-EGFP expression in the ventral ganglion, and axons within 100 µm of the ganglion. Scale bars are 20 µm. (**E**) Quantification of (**D**). **(F-G)** Retromer mutants exhibit accumulation of multiple EV and endosomally sorted cargoes. (**F**) Representative MaxIPs of EV (Syt4-EGFP) and non-EV (Tkv-mCherry, Syt1, CD8-mCherry) cargoes. (**G-I**) Quantification of Syt4-EGFP, Tkv-mCherry and Syt1 levels. Bar graphs show mean */− s.e.m.; dots show all data points representing individual NMJs. Scale bars are 5 µm. All measurements were normalized to presynaptic mean of their respective control (green line). **See Table 3 for detailed genotypes and statistical tests. See also Figure S2.**

To determine if retromer-dependent EV accumulation is specific to EV cargoes, we compared the effect of retromer loss on the accumulation of Synaptotagmin-4 (Syt4), an established EV cargo, and the BMP receptor Thickveins (Tkv), which localizes to presynaptic plasma membranes and endosomes and is not normally found in EVs (Deshpande et al., 2016; Korkut et al., 2013; O’Connor-Giles et al., 2008; Rodal et al., 2011). Endogenously tagged Syt4-EGFP exhibited increased pre- and-postsynaptic accumulation in *Vps35* mutants, as well as in mutants of *Vps26*, another component of the core retromer complex (**Fig 2F, G)**. Surprisingly, we also observed both presynaptic and postsynaptic punctate accumulation of Tkv-mCherry, similar to APP-EGFP and Syt4-EGFP (**Fig. 2F, H**). Thus, retromer loss leads to the general accumulation of multiple EV and endosomal cargoes and aberrant inclusion of non-EV cargoes into EVs. By contrast, two other neuronal transmembrane cargoes, Synaptotagmin-1 (Syt1) and CD8-mCherry, which do not undergo extensive endosomal trafficking at synapses, did not accumulate presynaptically and were not detected in EVs in *Vps35* mutants (**Fig 2F, I**) indicating that retromer may specifically sequester endosomally enriched transmembrane cargoes from the EV pathway.

Retromer is ubiquitously expressed, and retromer phenotypes related to *Drosophila* synaptic morphology and transmission depend on contributions from both neuronal and muscle retromer (Inoshita et al., 2017; Korolchuk et al., 2007). Indeed, we found that endogenously tagged Vps35-TagRFPt (Koles et al., 2015) localizes to both neurons and muscles (**Fig S2D**). Presynaptic Vps35-TagRFPt was faint and difficult to resolve from postsynaptic structures, so to further explore its localization, we expressed *Drosophila* Vps35-HA (Vps35-HA) in neurons using the GAL4-UAS system, and imaged synapses by SIM (**Fig S2E**). Vps35-HA was retained presynaptically, where it accumulated at the periphery of endosome-like structures and co-localized with a subset of APP-EGFP puncta. We tested the cell autonomy of retromer involvement in EV cargo trafficking by tissue-specific rescue of *Vps35* using the GAL4/UAS system. Neuronal restoration of Vps35-HA in the *Vps35* mutant background was sufficient to rescue both presynaptic and postsynaptic APP-EGFP to wild-type levels (**Fig 2A-C)**. Expressing Vps35-HA in neurons in a wild-type background had no effect on APP levels, indicating that rescue in the mutant was not confounded by Vps35 overexpression (**Fig S2F)**. Therefore, though Vps35 is present both pre and postsynaptically, and is required in both tissues to regulate synaptic growth and synaptic vesicle traffic (Inoshita et al., 2017; Korolchuk et al., 2007), presynaptic Vps35 is sufficient to restrict cargo trafficking into EVs and to prevent EV cargo accumulation.

*Vps35* mutants may have more EVs, or alternatively may package more cargo into the same number of EVs. To distinguish between these possibilities, we performed transmission electron microscopy on NMJs from control and *Vps35* mutant larvae. We noted a significant increase in postsynaptic 50-100 nm vesicles (consistent with the expected size for endosomally-derived exosomes) in the muscle SSR immediately surrounding the neuron (**Fig 3A,B**), suggesting that retromer mutant NMJs release more EVs.

**Figure 3.**
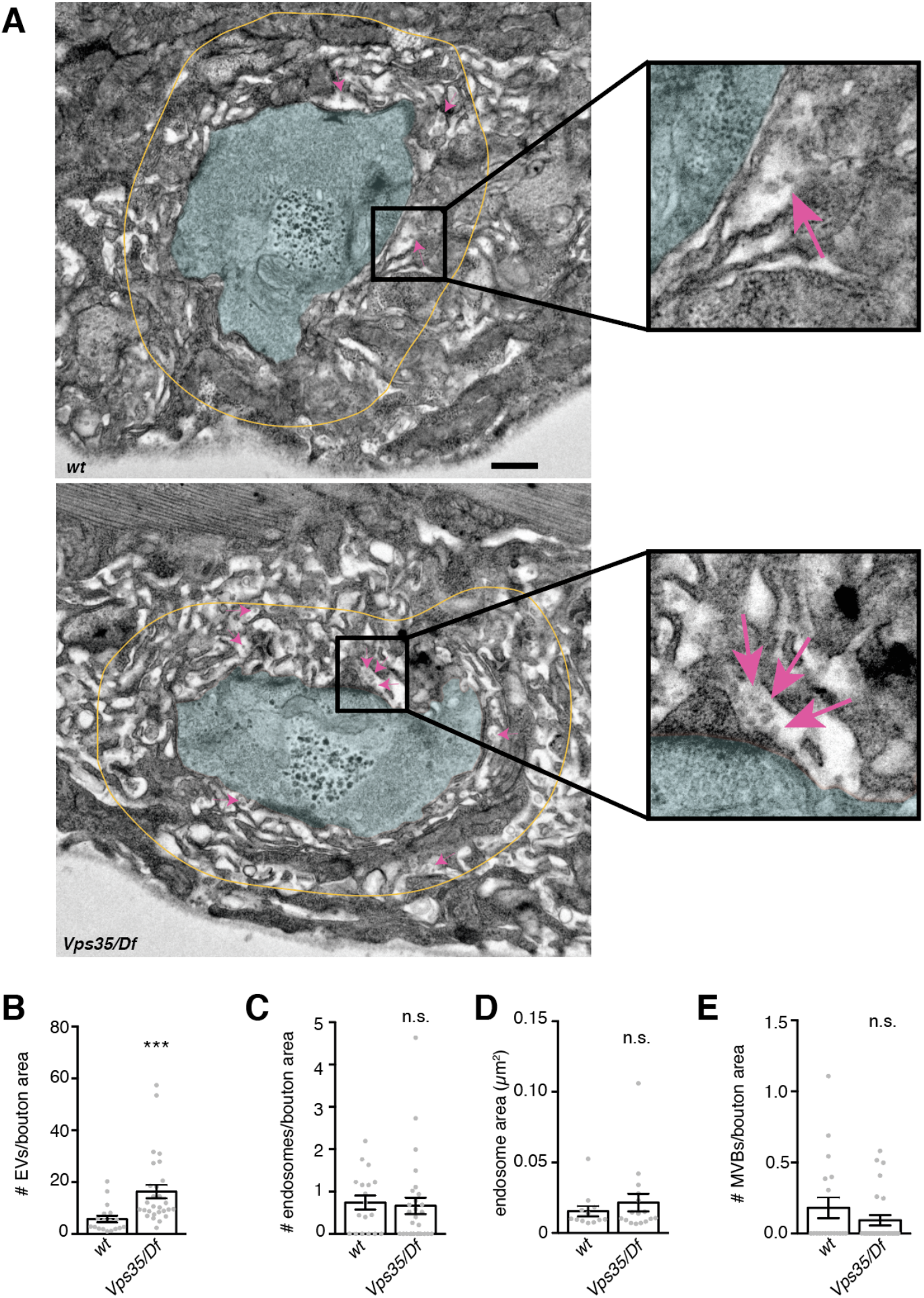
Retromer mutants exhibit more EV-sized vesicles in the perineuronal SSR, but do not exhibit changes in neuronal endosome profiles. **(A)** Transmission electron micrographs of thin sections showing muscle 6/7 boutons from larvae of the indicated genotype. Boutons are pseudocolored in cyan; yellow line indicates the perimeter of the region <1 µm from bouton plasma membrane, in which 40-100 nm vesicular structures were counted in the extracellular space. Magenta arrows indicate several examples of such EV-sized structures. Scale bar is 500 nm. Control is *w*^1118^. (**B**) *Vps35* mutants exhibit a greater number of 50-100 nm EV-sized vesicles, consistent with an increase in neuronally-derived EVs. However, neither the number nor size (**D**) of presynaptic endosomes (>100 nm vesicles) increases in a *Vps35* mutant. (**E**) Presynaptic MVBs are rare at both control and *Vps35* mutant synapses. Bar graphs show mean */− s.e.m.; each data point represents measurements derived from a single bouton. **See Table 3 for detailed genotypes and statistical tests.**

However, we did not observe a significant increase in presynaptic MVBs (defined as structures with intralumenal vesicles) or in the number or size of endosomes (defined as structures over 100 nm in diameter, to distinguish them from ~70 nm enlarged synaptic vesicles that are increased in *Vps35* mutants (Inoshita et al., 2017)) (**Fig 3C-E**). This result suggests that altered kinetics of MVB maturation or fusion with the plasma membrane (which would lead to MVB accumulation) are not a primary phenotype in *Vps35* mutants, and also that this step may not be rate-limiting for cargo and EV accumulation at the synapse.

### Retromer has separable functions in synaptic growth and EV traffic

One shared function of Appl and Vps35 at the *Drosophila* NMJ is to regulate the growth of synaptic arbors during larval development (Cassar and Kretzschmar, 2016; Korolchuk et al., 2007; Torroja et al., 1999) (**Fig 4A, B**). We considered the possibility that synaptic overgrowth in *Vps35* mutants is caused by accumulation of synaptic growth-relevant EV cargoes. To challenge this hypothesis, we asked if we could separate the functions of retromer in EV cargo traffic and synaptic growth by genetically manipulating retromer accessory factors. Retromer acts directly or indirectly with distinct combinations of sorting nexin (SNX) cofactors to direct specific trafficking events (McNally and Cullen, 2018; Wang et al., 2018). We measured synaptic growth in null mutants of *Snx27*, *Snx3*, and double mutants lacking heterodimeric ESCPE-1 (Endosomal SNX–BAR sorting complex for promoting exit 1) complex components *Snx1* and *Snx6* (Simonetti et al., 2019; Strutt et al., 2019; Zhang et al., 2011). We found that *Snx3* mutants exhibited significant synaptic overgrowth, while *Snx27* single and *Snx1*, *Snx6* double mutants did not (**Fig. 4A, B**). Retromer also interacts with the actin-regulating WASH complex (Chen et al., 2019b; Follett et al., 2014; McGough et al., 2014; Zavodszky et al., 2014), and a Parkinson’s Disease-associated mutant of human Vps35 (*Vps35*^D620N^) disrupts this association (Harbour et al., 2010; Jia et al., 2012; Seaman et al., 2013). Similar to previous findings (Inoshita et al., 2017; Malik et al., 2015), we found that knock-in of the analogous mutation (D628N) at the *Drosophila Vps35* locus (Koles et al., 2015) caused a robust synaptic overgrowth phenotype, similar to the *Vps35* null mutant (**Fig 4A, B**). These results suggest that SNX3 and WASH-associated functions of retromer are important for synaptic growth, while SNX27 and SNX1/6-associated functions are not.

**Figure 4.**
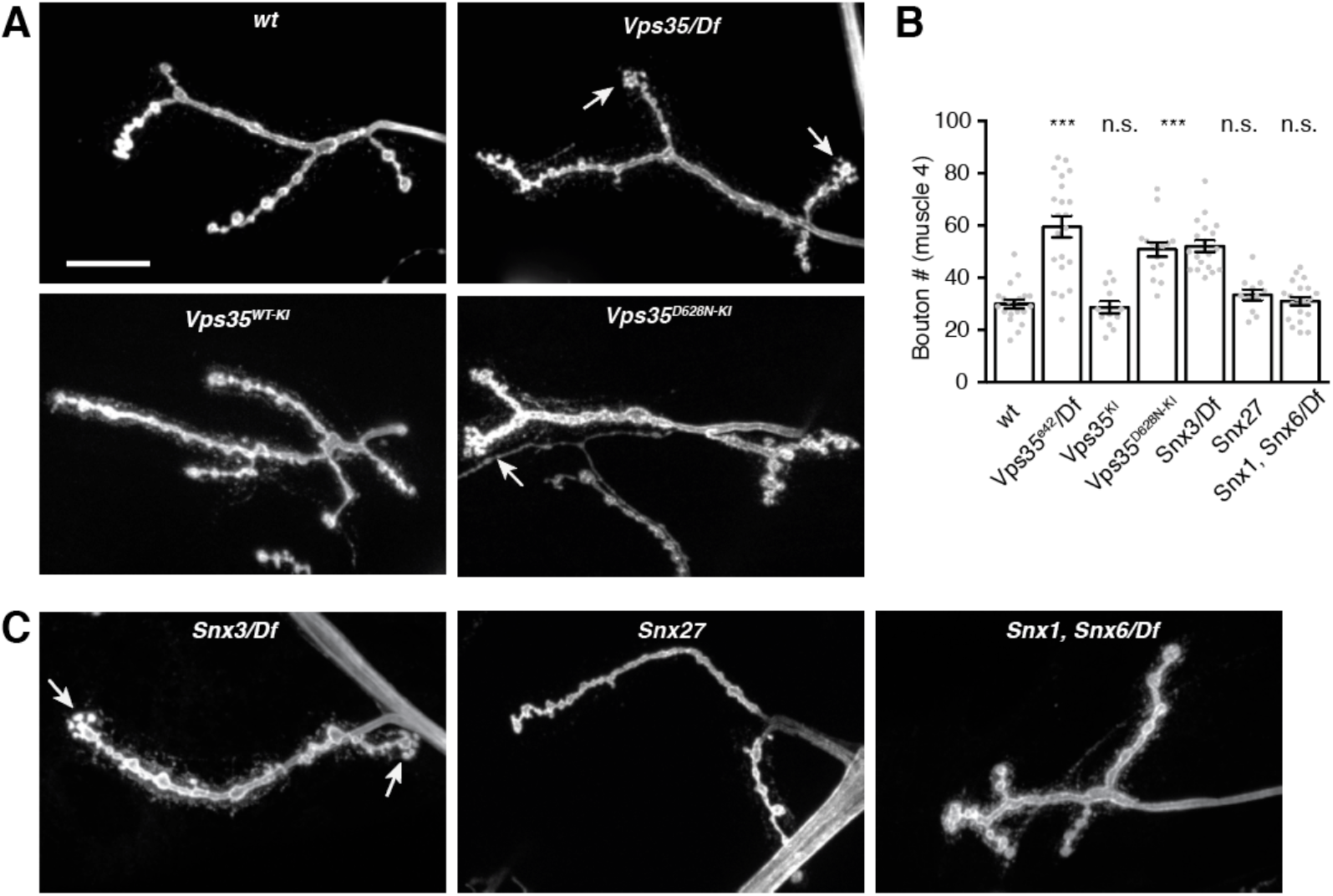
Control of synaptic growth depends on specific retromer-associated pathways. **(A,B)** *Vps35* null, *Snx3*, and *Vps35^D628N^* mutations cause an increase in bouton number at the NMJ, while mutations in retromer-associated *Snx1*, *Snx6* and *Snx27* have no effect. **(A)** MaxIPs of α-HRP-labeled muscle 4 NMJs in the indicated genotypes. Scale bar is 20 µm. Control is *w*^1118^. **(B)** Quantification of bouton number for muscle 4, segments A2 and A3.

We then asked if these separable functions of Vps35 in synaptic growth correlated with its EV cargo phenotypes. Remarkably, *Snx27*, *Snx3*, and *Vps35*^D628N^ mutants did not phenocopy the increased EV cargo levels of *Vps35* null mutants, while *Snx1/6* double mutants recapitulated the accumulation of presynaptic and postsynaptic APP-EGFP, though to a lesser extent than *Vps35* mutants (**Fig 5A-C**). These results demonstrate that synaptic growth and EV traffic regulation are distinct, non-correlated functions of retromer that can be genetically uncoupled, and that rely on separate components of the Vps35-associated machinery.

**Figure 5.**
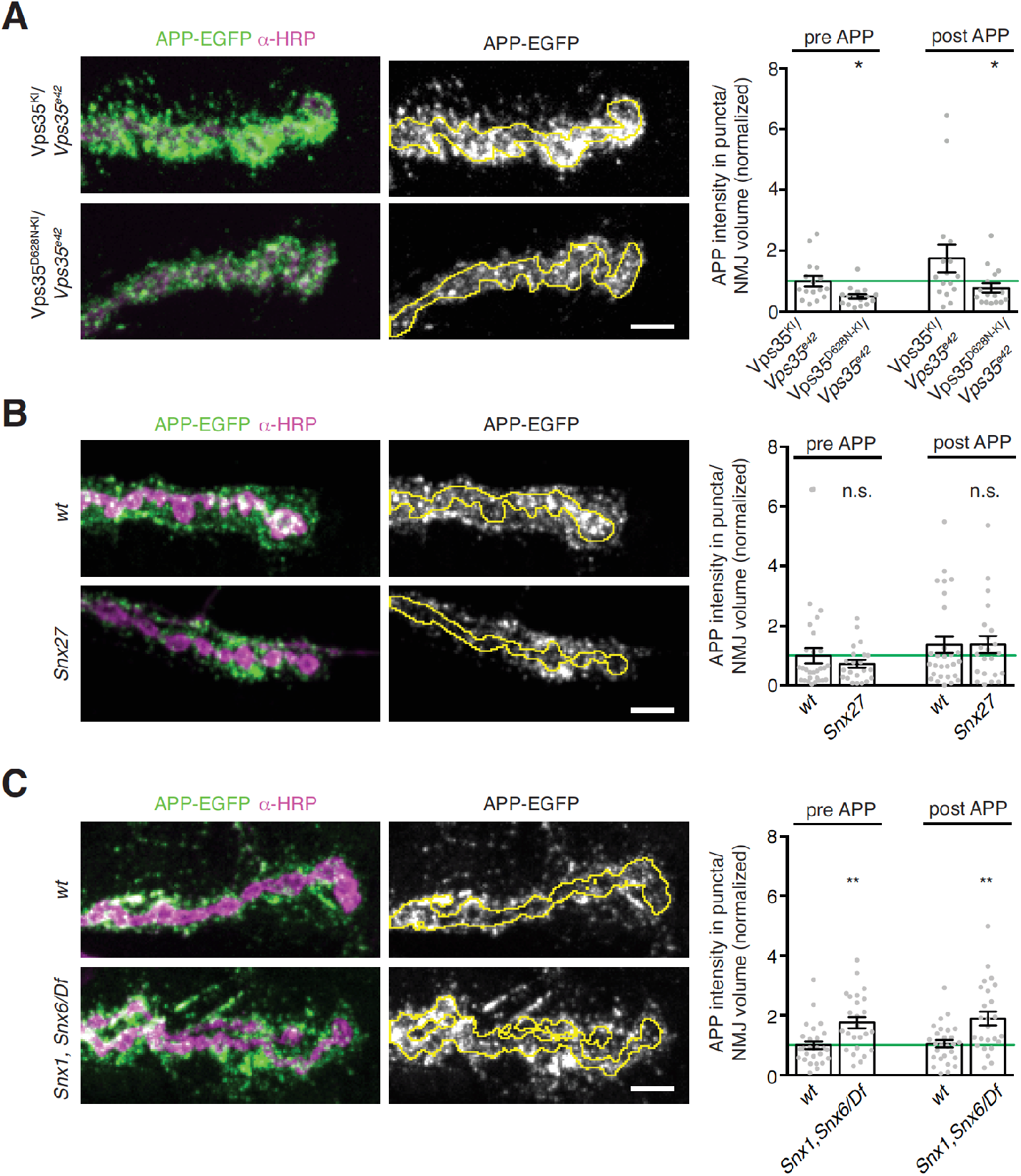
Control of EV Cargo levels depends on the Snx1/Snx6 ESCPE-1 complex but not *Vps35^D628^* or **Snx27**. (**A-C**) Representative MaxIPs and quantification of APP-EGFP levels at muscle 6/7 NMJs in the indicated genotypes Control for *Snx27* and *Snx1, Snx6* is GAL4^C155^. Each condition is normalized to the mean intensity of wild-type controls (green line). Bar graphs show mean */− s.e.m.; dots show all data points representing individual NMJs. Scale bars are 5 µm. **See Table 3 for detailed genotypes and statistical tests.**

### Lysosome dysfunction does not recapitulate EV phenotypes of Vps35 mutants

We next sought to determine the mechanisms by which loss of retromer results in increased levels of EV cargo. Since retromer is required for maintaining lysosomal hydrolase activity (Arighi et al., 2004; Cui et al., 2019; Seaman et al., 1998), one explanation for increased presynaptic and postsynaptic EV cargoes is general lysosomal dysfunction, leading to the accumulation and release of material that would otherwise be targeted for degradation (Becot et al., 2020). To address this hypothesis, we first examined the abundance of lysosomes at *Vps35* mutant NMJs. Spinster (Spin) is a putative Sphingosine-1-phosphate transporter that localizes to LysoTracker-positive lysosomes (Dermaut et al., 2005; Sweeney and Davis, 2002). We found that Spin-GFP intensity was significantly reduced at *Vps35* mutant NMJs (**Fig. S6A, B**), arguing against lysosomal expansion, and consistent with the absence of enlarged lysosomal profiles in our EM data. We next asked if lysosomes were functionally impaired in *Vps35* mutants by measuring the levels of Atg8-GFP, a marker of autophagosomes, which accumulate during lysosomal dysfunction. Instead, we observed a significant decrease in Atg8-GFP intensity in *Vps35* NMJs (**Fig. S6A, B**). Finally, staining with LysoTracker Deep Red, which labels acidic compartments, revealed fluorescence in the muscles that decreased in Vps35 mutants compared to controls (**Fig. S6C**). However, in presynaptic terminals we were unable to detect any fluorescence above background, in either control or *Vps35* mutants, again suggesting that acidic compartments are not amplified in *Vps35* mutant neurons. These results contrast with previous findings of increased LysoTracker and Atg8 fluorescence in *Drosophila Vps35* mutant fat body cells (Maruzs et al., 2015), and suggests that unlike in other tissues, Vps35 mutant synapses do not accumulate acidified endolysosomes or fail to degrade lysosomal cargoes.

We then asked if disruption of lysosomal function by orthogonal mechanisms affected EV cargo traffic similarly to the Vps35 mutant. We measured pre- and-postsynaptic Syt4-EGFP levels in mutants of *spin* and *deep orange* (*dor*/*Vps18*), which are both required for lysosome function at the NMJ (Dermaut et al., 2005; Fernandes et al., 2014; Nakano, 2019; Narayanan et al., 2000; Sweeney and Davis, 2002). In contrast to elevated Syt4-EGFP levels in *Vps35* mutant NMJs, pre- and postsynaptic Syt4-EGFP were significantly reduced in *spin* and *dor* mutant NMJs compared to controls (**Fig. S6D, E**). Finally, we pharmacologically disrupted lysosome function by raising wild-type larvae expressing Syt4-EGFP on food containing chloroquine, which blocks lysosome acidification (Gonzalez-Noriega et al., 1980). Chloroquine treatment caused a decrease in larval viability as well as enlarged presynaptic Syt4-EGFP puncta, indicating lysosome dysfunction. However, Syt4-EGFP puncta intensity did not change either pre- and-postsynaptically (**Fig. S6F, G**). Taken together, these results indicate that defects in the lysosomal pathway do not promote EV cargo accumulation, and that disruption of lysosomal degradation is unlikely to account for defects in neuronal EV cargo traffic in *Vps35* mutants.

### A Rab11-dependent recycling pathway maintains EV cargo levels

Our results indicate that generalized endolysosomal dysfunction is unlikely to account for Vps35 mutant EV phenotypes. Therefore, we next asked whether and how specific subtypes of endosomes were altered at synapses in *Vps35* mutants, by examining the localization and levels of Rab GTPases implicated in retromer traffic, using antibodies or tags knocked in to their endogenous loci (**Fig 6A**). GFP-Rab5, which is associated with early endosomes, showed a significant decrease in mean puncta intensity and number (but not size) in *Vps35* mutant NMJs compared to controls (**Fig. 6B-D**). By contrast YFP-Rab7, which is associated with late endosomes, showed a significant increase in intensity, puncta number, and puncta size in *Vps35* mutants, indicating an overabundance and enlargement of late endosomal compartments (**Fig. 6B-D**), consistent with the established role of retromer in attenuating Rab7 activity (Jimenez-Orgaz et al., 2017; Kvainickas et al., 2019; Seaman et al., 2018; Ye et al., 2020). Levels of α-Rab11 immunostaining, which marks the recycling endosome and post-Golgi secretory vesicles, remained unchanged, but Rab11 structures increased in size (but not number) (**Fig 6A, B, E-G)**. These results raised the possibility that one or several of these endosomal changes might underlie the retromer-dependent EV trafficking phenotypes, and thus define the site at which EV cargo fate is determined.

**Figure 6.**
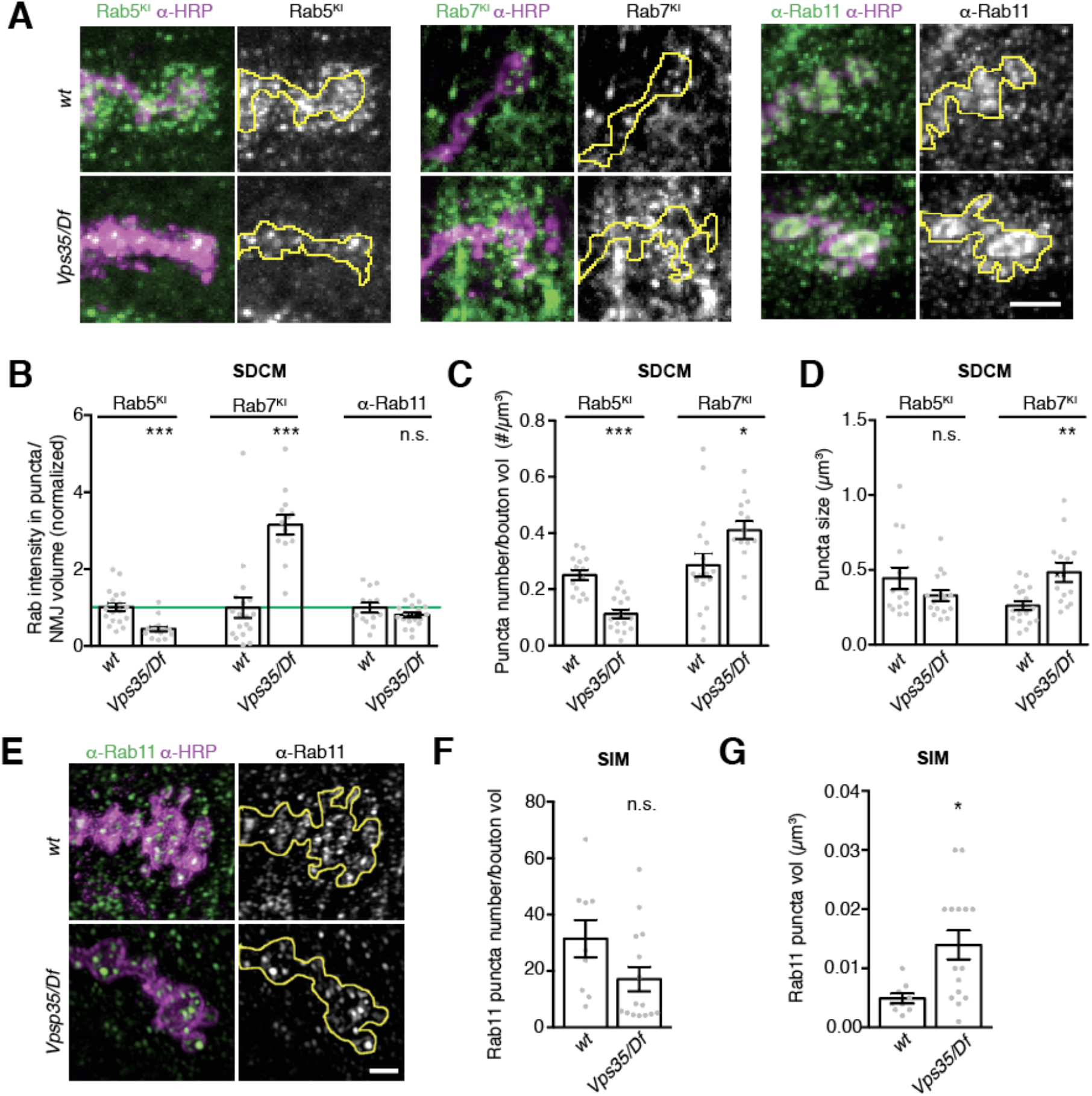
Endosome distribution in *Vps35* **mutants**. (**A**) MaxIPs of confocal images of NMJS from larvae expressing endogenously-tagged YFP-Rab7 or GFP-Rab5, or labeled with α-Rab11 antibodies, in control or *Vps35/Df* genotypes. Control is *w*^1118^. **(B)** Quantification of fluorescence intensity from spinning disc confocal microscopy (SDCM), normalized to mean intensity of wild-type control. **(C)** Rab5 and Rab7-positive puncta density by SDCM. **(D)** Rab5 and Rab7-positive size by SDCM. (**E**) SIM MaxIPs of NMJs from control or *Vps35/Df* genotypes, labeled with α-Rab11 antibody. Rab11 puncta were too dense to distinguish by SDCM, so we used SIM to analyze individual endosomes. By contrast, endogenous GFP-Rab5 and YFP-Rab7 intensities were insufficient for SIM imaging. Control is GAL4^C155^. (**F, G**) Quantification of puncta density and size from (**E**). Scale bars are 5 µm for SDCM, 2 µm for SIM. Bar graphs show mean */− s.e.m.; dots show all data points representing individual NMJs. **See also Figure S6.**

To ask which endosome subtype could be the site of retromer action in EV traffic, we separately perturbed their functions using neuron-specific expression of constitutively active (CA) or dominant negative (DN) forms of their resident Rab GTPases, and measured pre- and postsynaptic levels of endogenously tagged Syt4-EGFP. Overexpression of Rab5CA, which promotes homotypic fusion but blocks conversion to the Rab7-positive state (Rink et al., 2005; Stenmark et al., 1994), did not change presynaptic Syt4 levels, although Syt4 was redistributed into aberrant bright puncta. Notably, we observed decreased EVs in this condition. This suggests that trapping cargo in early endosomes inhibits EV release, and that the branchpoint at which retromer sorts cargo away from the EV pathway occurs downstream of Rab5. Overexpression of Rab5DN blocks homotypic endosome fusion and impairs maturation of the early endosome (Stenmark et al., 1994), so we used this manipulation to test if the decrease in Rab5 levels that we observed in our *Vps35* mutant could account for the increase in EV cargo levels. Indeed, we saw a twofold increase in presynaptic Syt4 levels, similar to *Vps35* mutants. However, unlike *Vps35* mutants, postsynaptic Syt4 was unaffected (**Fig 7A, B**). Thus, though inhibiting Rab5 function causes accumulation of EV cargoes in the presynaptic neuron, it does not affect release of these cargoes in EVs. This supports our previous conclusion that accumulation of cargo alone is not sufficient to drive EV sorting (**Fig. S2B**), and further suggests that the reduction in Rab5-positive compartments in *Vps35* mutants is unlikely to account for EV cargo sorting phenotypes.

**Figure 7.**
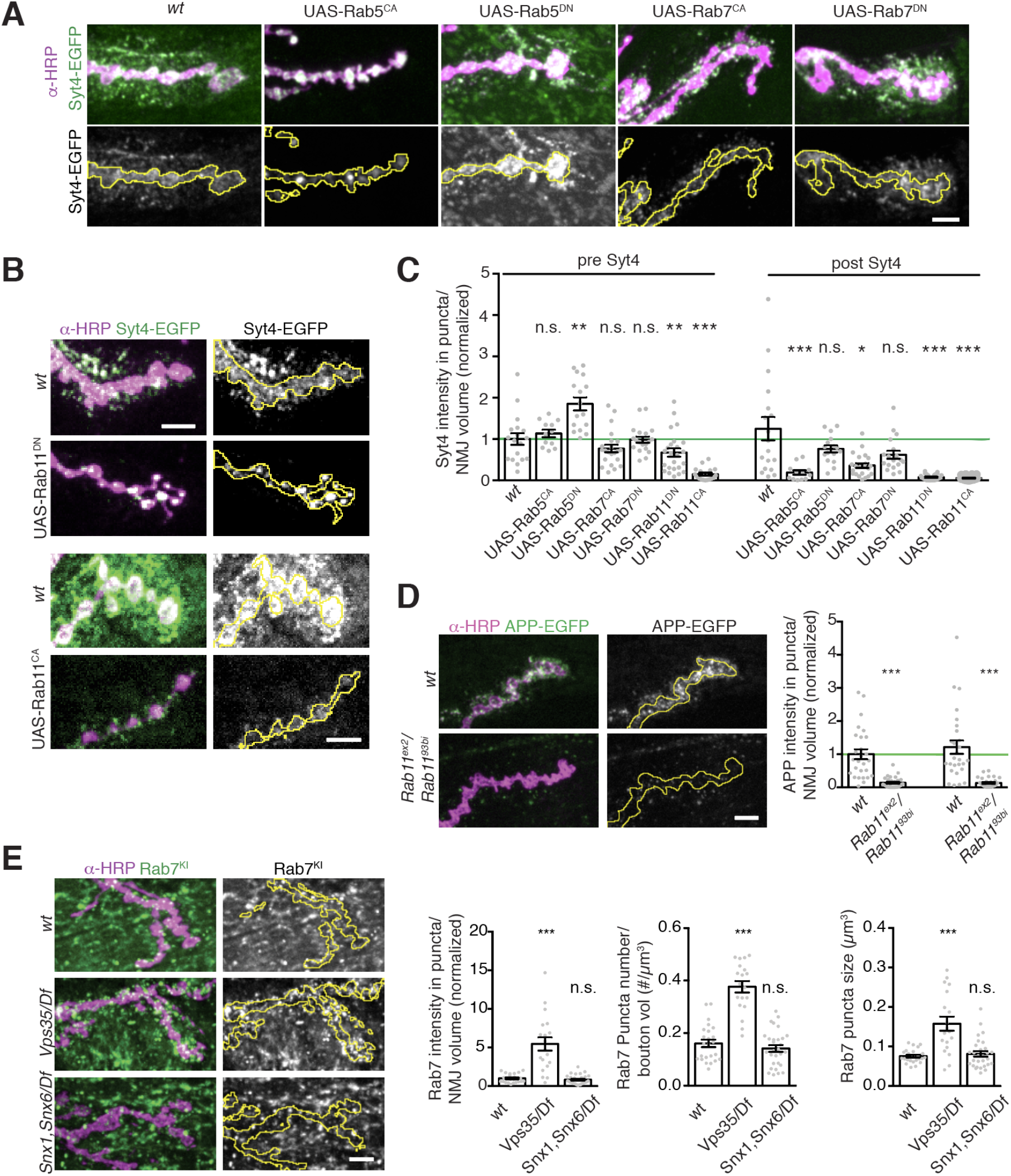
EV cargo accumulation depends on Rab11 activity. (**A-B**) MaxIPs and quantification of NMJs from larvae expressing endogenously-tagged Syt4-EGFP with neuronally-driven (GAL4^C380^) dominant negative (DN) or constitutively active (CA) Rab GTPase transgenes. Control is GAL4^C380^. **(B)** Quantification of (**A, B**), normalized to presynaptic mean intensity of wild-type control. All Rab5 and Rab7 manipulations are normalized to the control shown on the graph; Rab11 manipulations were performed using different imaging parameters and are normalized to their own controls. (**D** (Left) MaxIPs of NMJs expressing APP-EGFP in control or Rab11 mutant backgrounds. (Right) quantification, normalized to mean of presynaptic control. Control dataset (GAL4^C155^) is identical to **Fig. 2B**. (**E**) (Left) MaxIPs of NMJs expressing YFP-Rab7 in control or *Snx1, Snx6* mutant backgrounds. (Right) Quantification of presynaptic Rab7 intensity and endosome number and size. Scale bars are 5 µm. Bar graphs show mean */− s.e.m.; dots show all data points representing individual NMJs.

Manipulations of Rab7 had a milder influence overall on Syt4 trafficking compared to Rab5. Overexpression of Rab7^CA^ caused a modest decrease in EV Syt4 (**Fig 7A, B)**. Rab7^CA^ increases the rate by which the late endosome merges with the lysosome (Meresse et al., 1995), so these phenotypes may reflect accelerated degradation of EV cargoes. Rab7^DN^ overexpression, which inhibits recruitment of Vps35 to late endosomes (Rojas et al., 2008; Seaman et al., 2009) and prevents the Rab5-to-Rab7 switch that is a hallmark of endosome maturation (Rink et al., 2005), had no effect on pre-or-postsynaptic Syt4 levels (**Fig 7A, B**). Finally, we examined the Rab7 compartment in *Snx1*, *Snx6* ESCPE-1 double mutants, which exhibit increased EV cargo levels (**Fig. 5**). Rab7-positive endosome intensity, size and number were not significantly different from wild type in ESCPE-1 mutants (**Fig. 7E**). Together, these data argue that the accumulation of Rab7 and Rab7-associated compartments in *Vps35* mutant is unlikely to be necessary for or cause the EV cargo trafficking defect, and that synaptic EV cargo sorting does not depend on recruitment of Vps35 to late endosomes by Rab7.

Finally, we examined the effects of manipulating Rab11. Dominant negative and loss-of-function alleles of Rab11 block recycling of endosome- and Golgi-derived carriers to the plasma membrane (Ullrich et al., 1996), and have previously been shown to reduce EV levels of the Transferrin receptor (TfR), Evi, Syt4, and Arc (Ashley et al., 2018; Beckett et al., 2013; Koles et al., 2012; Korkut et al., 2013; Savina et al., 2002). These findings led to the hypothesis that Rab11 is involved in MVB fusion with the plasma membrane, as well as the prediction that EV cargoes should accumulate presynaptically in Rab11 mutants, though this has not yet been directly tested. Surprisingly, opposite to this prediction, we found a strong decrease of both pre- and post-synaptic Syt4-EGFP upon Rab11^DN^ expression (**Fig. 7B, C**), and of APP-EGFP in a hypomorphic *rab11* loss-of-function allelic combination (**Fig 7E, F**). By contrast, we previously found that Rab11^DN^ has no effect on presynaptic levels of the non-EV cargo Tkv (Deshpande and Rodal, 2016), indicating that this depletion is specific to EV cargo. Constitutively active Rab11 (Rab11^CA^) is predicted to enlarge the recycling endosome and also disrupts recycling activity (Ullrich et al., 1996; Wilcke et al., 2000). We found that neuronal expression of Rab11^CA^ caused a dramatic defect in neuronal morphology including reduced α-HRP staining, synaptic bouton size, and overall arbor size (as previously described (Akbergenova and Littleton, 2017)). With the caveat that this extreme phenotype may cause pleiotropic defects in membrane traffic, Syt4-EGFP levels were also dramatically reduced both presynaptically and postsynaptically in Rab11^CA^-expressing animals. Since Rab11 manipulations caused EV phenotypes opposite from Vps35 mutants, we hypothesized that Rab11 may regulate EV cargo levels by counteracting the Vps35 pathway. We therefore further investigated the role of Rab11 compartments in the *Vps35* mutant phenotype.

### APP accumulates in a Rab11-dependent compartment in Vps35 *mutants*

Depending on the specific cell type and cargo, loss of retromer causes differential effects on cargo flux through the endolysosomal pathway, and can result in altered cargo levels either in early endosomes, late endosomes, or on the plasma membrane. (Chen et al., 2013; Hussain et al., 2014; Jimenez-Orgaz et al., 2017; Loo et al., 2014; Pocha et al., 2011; Steinberg et al., 2013; Strutt et al., 2019; Tian et al., 2015; Vazquez-Sanchez et al., 2018; Wang et al., 2014; Wang et al., 2013; Zhou et al., 2011). One hypothesis for how retromer restricts pre- and-postsynaptic accumulation of APP or Syt4 in neurons is by preventing cargo accumulation at a stage in the endosomal pathway that is permissive for EV release in this cell type. Our genetic data indicate that this may be the Rab11 compartment, so we used SIM to directly ask if APP traffics through presynaptic Rab11-positive compartments, and if this is altered in *Vps35* mutants. In control animals, APP-EGFP and Rab11 partially co-localized (**Fig. 8A**), and in *Vps35* mutants, we found an increased fraction of Rab11 signal in APP-positive compartments (**Fig. 8A-C**). These data suggest that when EV cargo is retained in a recycling regime, it is more likely to be incorporated into EVs.

**Figure 8.**
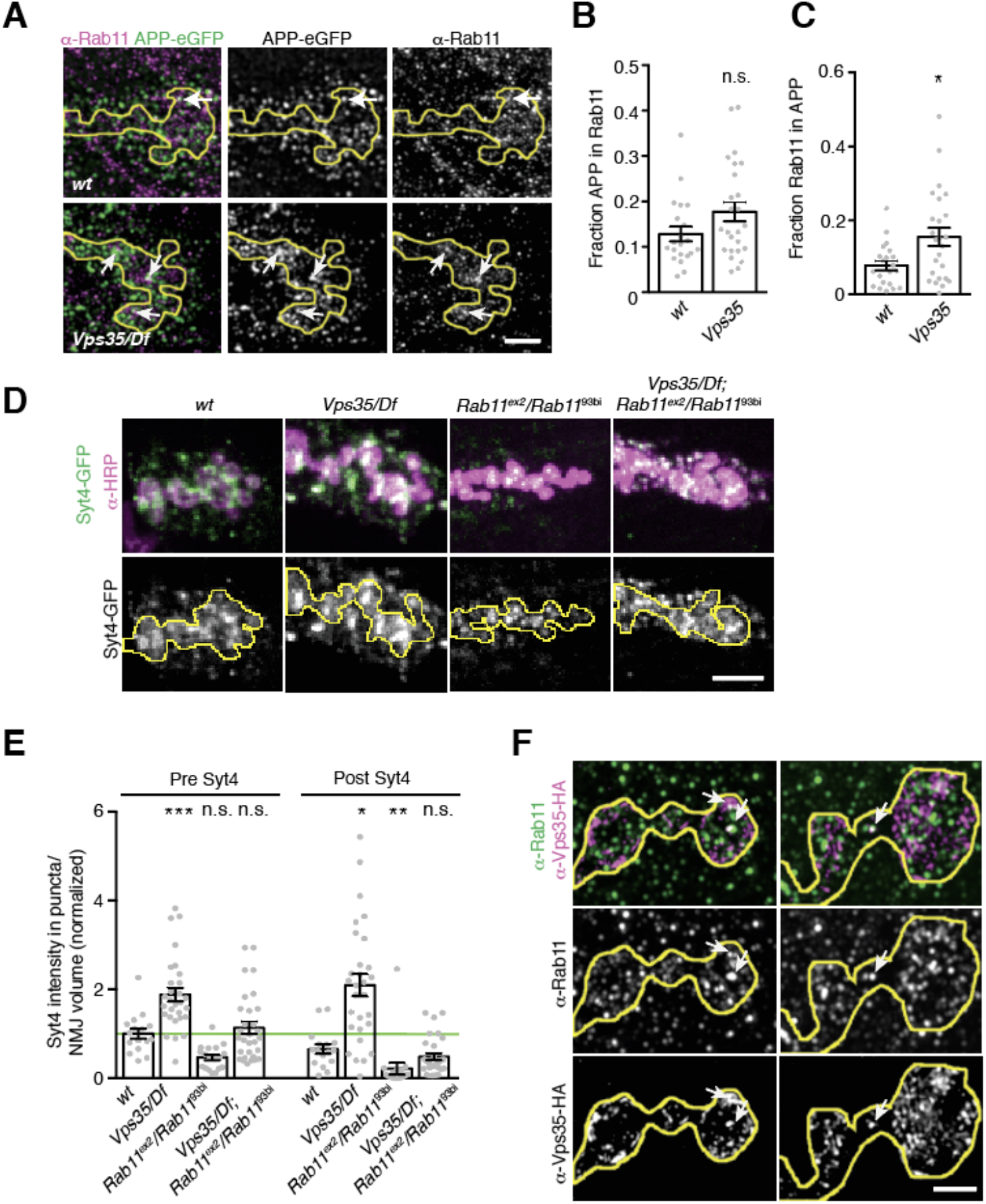
EV cargo accumulates in a Rab11-positive and - dependent compartment in *Vps35* **mutants**. (**A**) MaxIPs of SIM images showing control and *Vps35* larvae expressing GAL4^C155^-driven UAS-APP-EGFP and stained with α-Rab11. White arrows indicate instances of colocalization. Scale bar is 2 µm. Control is GAL4^C155^. (**B,C**). Quantification of Mander’s coefficients from (**A**). *Vps35* NMJs exhibit increased fraction of presynaptic Rab11 overlapping with APP. (**D,E**) Rab11 and Vps35 act in opposing pathways. (**D**) MaxIPs of NMJS from larvae expressing endogenously-tagged Syt4-EGFP in the indicated genotypes. Control is *w*^1118^. (**E**) Quantification of (**D**), normalized to presynaptic mean intensity of wild-type control. (**F**) Vps35 and Rab11 colocalize on a small subset of endosomes. MaxIP SIM images from NMJs expressing neuronally-driven Vps35-HA immunolabeled for α-HA and α-Rab11. Arrows indicate examples of structures that are positive for both Vps35-HA and Rab11. Scale bar is 2 µm. Bar graphs show mean */− s.e.m.; dots show all data points representing individual NMJs.

We then asked whether aberrant sorting of EV cargo through this Rab11-positive compartment in *Vps35* mutants caused its accumulation, by testing if levels could be restored by rebalancing the opposing activities of Rab11 and Vps35 in *Vps35*; *rab11* double hypomorphic mutants. Strikingly, loss of *rab11* function in the *Vps35* mutant restored both pre- and post-synaptic Syt4 to wild type levels (**Fig. 8D, E**). These results suggest that accumulation of Syt4 in a Rab11-dependent compartment (possibly in the Rab11-positive compartment itself) in Vps35 mutants is causative of EV cargo accumulation and release. Finally, to ask whether Rab11 and Vps35 might be acting on the same compartment, we performed SIM and found Rab11-Vps35 positive puncta in all NMJs we examined (n=8, **Fig 8F**).

## DISCUSSION

Here, we find that EVs represent a major trafficking trajectory for neuronal APP *in vivo*, and that this pathway is restricted by the neuron-autonomous activity of retromer. Rather than playing a general role in maintaining overall endolysosomal homeostasis, we find that retromer has separable roles in neurons and specifically restricts traffic within the EV pathway by regulating accumulation of cargoes in Rab11-dependent endosomes. Our results indicate that retromer and Rab11 balance traffic of AD-relevant cargoes such as APP into the EV pathway, and may therefore be critical for disease propagation in the brain.

### Neuronal retromer has multiple, separable functions

Our finding of EV cargo traffic as a new function for neuronal retromer adds to its previously described roles in synaptic morphogenesis, synaptic transmission, synaptic vesicle loading and recycling, and AMPA receptor traffic (Bhalla et al., 2012; Choy et al., 2014; Inoshita et al., 2017; Korolchuk et al., 2007; Munsie et al., 2015; Temkin et al., 2017; Tian et al., 2015; Vazquez-Sanchez et al., 2018; Wu et al., 2017). Interestingly, non-EV associated phenotypes of neuronal retromer, including photoreceptor degeneration and synaptic transmission defects, are opposed by Rab11, similar to our findings for the EV phenotype, suggesting that balance between these pathways is of general importance in neurons (Inoshita et al., 2017; Satoh et al., 2005; Wang et al., 2014). However, our data show that the role of retromer in EV traffic is separable from its other functions. Compared to synaptic transmission and synaptic growth defects, which require both neuronal and muscle retromer (Inoshita et al., 2017; Korolchuk et al., 2007), EV cargo regulation depends strictly on presynaptic retromer. Further, our data show that synaptic growth, Rab7 endosome maturation and EV defects are genetically separable: *Snx3* and *Vps35*^D628N^ mutants phenocopy *Vps35* null mutants for synaptic growth (likely via excess BMP signaling (Korolchuk et al. 2007)) but not for EV traffic, while ESCPE-1 mutants recapitulate EV but not synaptic growth or Rab7 turnover phenotypes of Vps35.

Given these diverse phenotypes, it will be critical to determine which of these many cellular functions of retromer underlie neurological disease symptoms. Indeed, acute depletion of Vps35 in hippocampal neurons leads only to endosomal trafficking defects, but not to synaptic vesicle cycling or morphogenesis defects that are observed upon chronic retromer depletion (Vazquez-Sanchez et al., 2018). While chronic phenotypes remain interesting in light of retromer depletion in AD, these data indicate that some retromer-associated neuronal defects may be indirect or even compensatory. Overall, the diverse cellular roles of neuronal retromer may differentially contribute to neurological disease, and individually manipulating these functions via distinct retromer-associated molecular pathways may improve therapeutic strategies for disease treatment (Berman et al., 2015; Mecozzi et al., 2014).

### Retromer and Rab11-dependent mechanisms of EV traffic in neurons

Our data reinforce the important concept that effects of retromer perturbation on the endosomal system are both cargo and cell type-specific, and highlight the importance of studying the cell biological functions of retromer in the appropriate cell type *in vivo*. The canonical role of retromer is to rescue cargoes from lysosomal degradation and to redirect them either to endosome-to-Golgi (e.g. the cation-independent mannose 6-phosphate receptor (CI-MPR) or endosome-to-plasma membrane (e.g. β-adrenergic receptor) recycling routes; in the absence of retromer both these classes of cargoes are misrouted to the lysosome and degraded (Arighi et al., 2004; Cui et al., 2019; Seaman et al., 1998; Steinberg et al., 2013; Temkin et al., 2011). By contrast, we observe strong accumulation of cargoes in *Vps35*-deficient *Drosophila* neurons, similar to observations in *Vps35*-depleted hippocampal neurons (Bhalla et al., 2012), or in secretory cells such as the *Drosophila* salivary gland (Neuman et al., 2020). A prevailing hypothesis that could account for cargo accumulation in endosomes and EVs is retromer-dependent lysosome dysfunction (Maruzs et al., 2015; Miura et al., 2014; Wang et al., 2014). Indeed, lysosomal damage increases APP-CTFs in secreted EVs from neuroblastoma cells (Miranda et al., 2018; Vingtdeux et al., 2007). However, in our *in vivo* system, we did not observe the functional or ultrastructural indications of lysosome defects that are seen in other tissues. Further, we did not phenocopy the *Vps35* mutant phenotype by manipulating late endosome or lysosome function, similar to findings in the salivary gland salivary gland (Neuman et al., 2020), suggesting it is not sufficient to account for the observed cargo increase (though we cannot exclude that it plays a contributing role).

Our data instead suggest that in retromer mutants, neuronal endosomal cargoes are diverted to the EV pathway, rather than being shunted to lysosomal degradation as is seen in other cell types. This could be beneficial, since EV recipient cells (in this case muscles and glia) may be better suited for subsequent cargo degradation than the donor neuron synaptic terminal, which may have limited lysosomal capabilities (Eitan et al., 2016; Ferguson, 2018; French et al., 2017; Fuentes-Medel et al., 2009). According to this model, in presynaptic nerve terminals, MVBs would preferentially fuse with the plasma membrane rather than with lysosomes. Why then do we not observe excess MVBs by electron microscopy in retromer mutants, and why do EV cargoes accumulate in Rab11-positive compartments?

We propose that EV traffic is not rate-limited by MVB-plasma membrane fusion (which may occur rapidly via the abundant factor Hsp90 (Lauwers et al., 2018)), but instead is kinetically restricted by loading of early endosomes and MVBs with EV cargo through a Rab11-plasma membrane recycling pathway (**Fig 9**). In support of conservation of recycling flux for EV traffic in other neuronal cell types, EV-directed p75-neurotrophin receptor accumulates in Rab11-positive compartments in both PC12 cells and sympathetic neurons (Escudero et al., 2014). Rab11 has previously implicated in EV traffic, and suggested to be involved in MVB-plasma membrane fusion (Escudero et al., 2014; Fan et al., 2019; Koles et al., 2012; Korkut et al., 2013; Savina et al., 2002). Our results instead show that the primary cause of EV trafficking defects in *rab11* mutants is cargo depletion in the donor cell (as has previously been shown for the EV cargo Arc (Ashley et al., 2018)). We hypothesize that Rab11 maintains a cellular pool of EV cargo that cycles between early endosomes, recycling endosomes, and the plasma membrane; this recycling pool of cargo is protected from lysosomal degradation, and is incorporated into MVBs destined to become EVs (EV-MVBs). We also cannot exclude the possibility that EV cargo is released during this recycling itinerary by direct budding from the plasma membrane in microvesicles. An alternative hypothesis is that the recycling pathway directly loads cargo into a Rab11-positive MVB-forming endosome, as has been proposed for EV-releasing *Drosophila* male accessory gland secondary cells (Corrigan et al., 2014; Fan et al., 2019; Marie et al., 2020).

**Figure 9.**
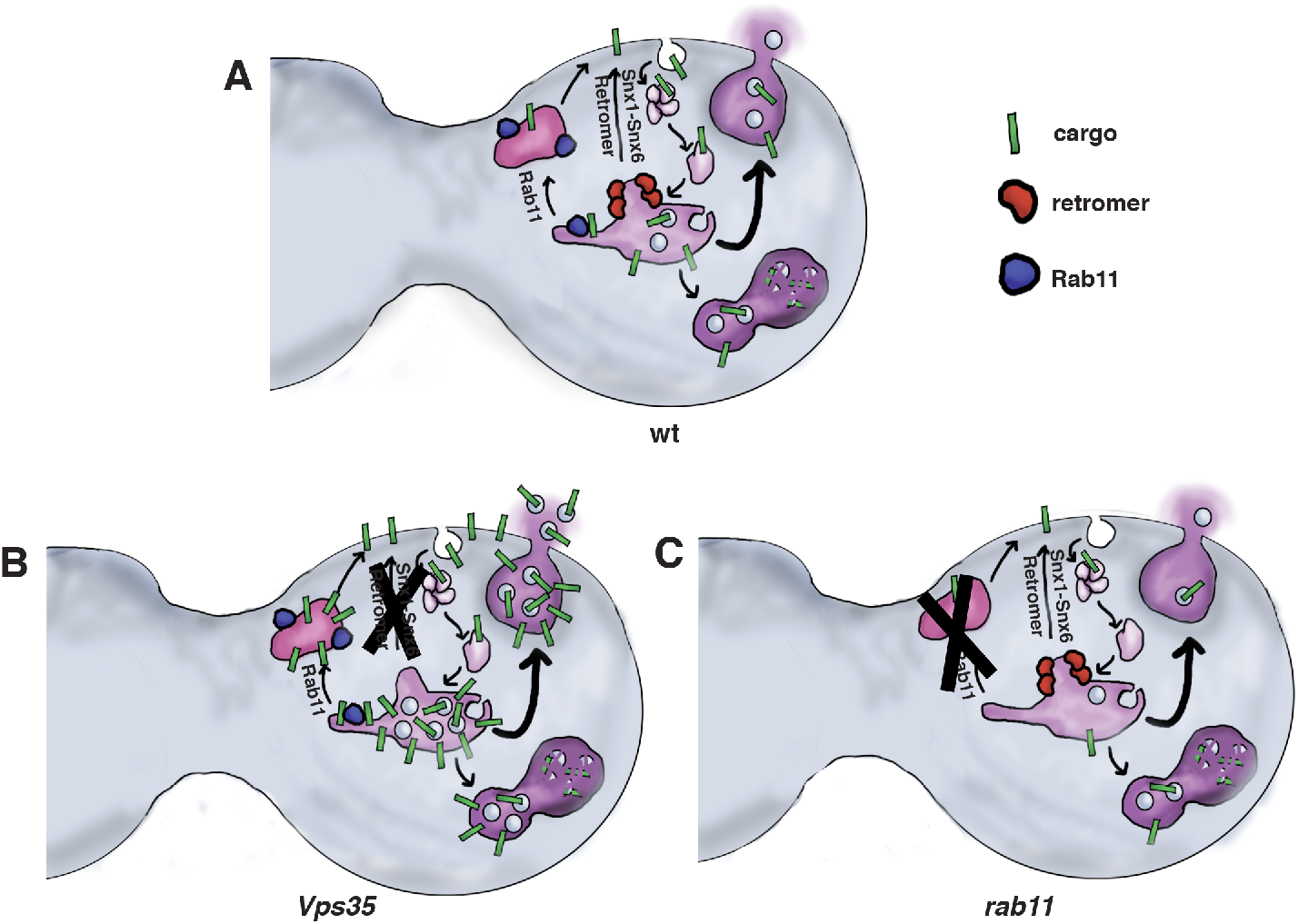
Model for functions of Vps35 and Rab11 in neuronal EV cargo traffic. **(A)**At wild-type synapses Rab11 maintains a rapidly-recycling pool of EV cargo between the sorting/early endosome, the recycling endosome, and the plasma membrane. MVBs in this pathway preferentially fuse with the plasma membrane and release their contents as ILVs. Retromer removes cargo from the sorting endosome and delivers it to other destinations such as the plasma membrane or the trans-Golgi network. (**B**) In a *Vps35* mutant, cargo is retained in the recycling pathway. Increased levels of cargo on the endosome membrane may drive ILV formation, resulting in EV secretion and pre- and -postsynaptic cargo accumulation. (**C**) In a *rab11* mutant, the recycling pool of cargo is not maintained. Cargo may proceed to the lysosome for degradation and is depleted from the synapse.

Retromer, working in concert with ESCPE-1, may oppose EV formation in one of two non-exclusive ways: First, it might retrieve or protect cargo from slow Rab11-dependent trafficking through the recycling endosome (for example via fast and direct recycling to the plasma membrane (Naslavsky and Caplan, 2018; Steinberg et al., 2013)), either by specifically interacting with cargo itself (Simonetti et al., 2017) or via carriers such as SorLA (Fjorback et al., 2012; Nielsen et al., 2007)). Our identification of ESCPE-1 as a relevant retromer cofactor will be informative in future studies of EV cargo sorting determinants, given recent insights into sorting motifs recognized by the SNX1-SNX6 heterodimer (Simonetti et al., 2019; Yong et al., 2020). In support of this direct sorting hypothesis, we found that Rab11 and Vps35 co-localize at the synapse, suggesting that they may sort cargo from the same compartment. Second, retromer and ESCPE-1 may non-specifically restrict MVB formation from early endosomes (by creating endosomal domains protected from ESCRT-mediated ILV formation (Derivery et al., 2012; Norris et al., 2017), or by limiting ESCRT activity-promoting cargo accumulation on endosomes (Baietti et al., 2012)). In support of this latter hypothesis, we found in *Vps35* mutants that the non-EV cargo Tkv was aberrantly trafficked into EVs, and that there was an overall increase in EV-like postsynaptic structures. Importantly, our data suggest that retromer’s role in promoting Rab7 turnover and late endosome maturation is unlikely to be involved in EV traffic at this synapse, though Rab7 has been implicated in EV release in other neuronal cell types (Song et al., 2016).

### Role of the retromer-Rab11 EV pathway in neuronal function and disease

Our results have significant implications for understanding the functional and pathological consequences arising from loss of Vps35. We found that the Parkinson’s Disease-modeling *Vps35*^D628N^ mutant does not recapitulate the EV trafficking defects arising from deletion of *Vps35*, suggesting that EVs may not play a primary role in patients with this mutation. Instead, our results in the *Vps35* deletion mutant may relate more directly to AD, which is associated with reduced overall levels of retromer based on genetic and pathological data from patients (Small et al., 2005; Vardarajan et al., 2012), and findings that loss of *Vps35* enhances pathology in animal models of AD (Li et al., 2019; Muhammad et al., 2008; Wen et al., 2011). The specific role of retromer in APP traffic has remained unclear, because of differing results in various cell types regarding trafficking itineraries and points of intersection for APP and its processing proteases (Bhalla et al., 2012; Brodin and Shupliakov, 2018; Choy et al., 2012; Cuartero et al., 2012; Das et al., 2013; Das et al., 2016; Fjorback et al., 2012; Li et al., 2012; Sullivan et al., 2011; Tan and Gleeson, 2019; Toh et al., 2017; Vieira et al., 2010; Wang et al., 2012a; Wen et al., 2011). In neurons, loss of retromer increases the co-localization of APP and BACE1 in early and late endosomes, resulting in higher Aβ levels (Bhalla et al., 2012; Toh et al., 2017; Wang et al., 2012a; Wen et al., 2011). Our results now extend the role of neuronal retromer to the EV pathway, and show that retromer loss promotes traffic of endosomally derived APP CTFs into synaptic EVs, potentially driving intercellular propagation of Aβ as well as other disease-relevant EV cargoes such as tau (Kanmert et al., 2015).

Conversely, previous data show that loss of Rab11 suppresses AD-related phenotypes. Rab11 variants have been associated with late-onset AD risk, loss of Rab11 leads to decreased Aβ, and activation of Rab11 leads to increased Aβ (Li et al., 2012; Udayar et al., 2013). These effects were previously interpreted in the context of functions of Rab11 in intracellular traffic (e.g. by regulating the endocytosis of Aβ as well as the recycling and axonal transport of BACE1 (Buggia-Prevot et al., 2014; Li et al., 2012; Udayar et al., 2013)), but should be reconsidered in light of the unexpected role we found for Rab11 in regulating synaptic EV cargo levels. Overall, our results indicate that in addition to the previously described opposing effects of retromer and Rab11 on intracellular processing of APP, the phenotypes previously observed for these perturbations in cellular and animal models, as well as human disease, may also arise from their opposing roles in the neuronal EV pathway.

What then is the physiological function of the retromer-dependent EV pathway? In the context of AD, and more generally for neuronal proteostasis, our results suggest that an important role of neuronal retromer-Rab11 EV pathways is to control delivery of neuronal cargo to recipient cells for disposal, as an alternative to intraneuronal lysosomal degradation. However, it is also important to consider increasing evidence that neuronal EV cargoes play functional roles in recipient cells, driving both structural and functional plasticity as well as morphogen signaling, in an activity-dependent fashion (Ashley et al., 2018; Coulter et al., 2018; Faure et al., 2006; Gong et al., 2016; Korkut et al., 2009; Korkut et al., 2013; Pastuzyn et al., 2018). In the future, it will be exciting to test whether neurons control these functional processes by directly tuning EV traffic via retromer or Rab11, lending these trafficking pathways interesting new roles beyond general synaptic proteostasis.

## MATERIALS AND METHODS

### *Drosophila* strains and methods

*Drosophila Tkv* and human APP^695^ were cloned into pBI-UASc-gateway-EGFP or pBI-UASc-gateway-mCherry, which were generated from Gateway-AttB vectors (Wang et al., 2012b), and injected into flies (Genetic Services Inc. Cambridge, MA), using ϕc381 recombinase at the Attp40 locus (Ni et al., 2008). The endogenous Syt4 locus was tagged with a tissue-specific convertible TagRFPt to EGFP tag (T-STEP (Koles et al., 2015)), followed by germline conversion to EGFP to create an EGFP knockin. Briefly, the following genomic sequence was used to target DNA cleavage by Cas9: TGAACGAGTAGGGGAGGGGC (at location 3R:7269302-7269321), with the 5’ T converted to the obligatory G, thus yielding the 20 bp sequence GGAACGAGTAGGGGAGGGGC. The T-STEP cassette was flanked with 5’ homology (3R: 7267814-7269309) and 3’ homology (3R:7269313-7270955) arms, and gene targeting was carried out as described ((Koles et al., 2015), based on (Chen et al., 2015)). Syt4-TSTEP flies were crossed to bam-Gal4;;UAS-Rippase::PEST @attP2 flies to isolate germline ripouts of the TagRFPt cassette, leaving a C-terminal EGFP knockin with the following linker (Syt4 sequence in lowercase, EGFP in italics, Prescission protease site underlined): prrqiaewhklneSYALMKEYVILSSSSGSSLEVLFQGPGSGSGS*MVSK GEELFTGV.* All other fly stocks and sources are described in **Table 1**.

*Drosophila* larvae were cultured using standard media at controlled density at 25º for all experiments. Wandering 3rd instar larvae were dissected in calcium-free HL3.1 saline (Feng et al., 2004) and fixed in HL3.1 containing 4% formaldehyde before antibody staining. Male larvae were used for all experiments, except for chloroquine treatment (Fig **S6F-G)**and *Vps35 rab11* genetic interaction experiments (**Fig 8D-E**), for which both male and female larvae were used. For Chloroquine treatment, Formula 4-24 Instant *Drosophila* Medium (Carolina Biological Supply Company, Burlington, NC) was prepared at a 1:2 ratio with 0-8 mM Chloroquine diluted in water, and allowed to solidify at room temperature. 4 male and 4 female larvae homozygous for Syt4-EGFP were added to each vial, and vials were incubated for up to 9 days at 25ºC.

### Immunohistochemistry and Imaging

For primary antibody information and concentrations, see **Table 2.** FluoTag-X4 α-GFP nanobodies (Nanotag Biotechnologies) were used to amplify EGFP signal for SIM imaging only; all other EGFP imaging reflects endogenous EGFP fluorescence. All primary and secondary incubations were conducted in PBS with 0.5% TritonX-100 and 2% (w/v) BSA and 2% (v/v) normal goat serum, and all washes in PBS with 0.5% TritonX-100. α-HRP antibodies and secondary antibodies for imaging were conjugated to Dylight 488, Rhodamine Red-X, or Alexa 647 (Jackson Immunoresearch) and were used at a concentration of 1:250 (Alexa 647) or 1:100 (Dylight 488, Rhodamine Red-X). For LysoTracker staining, fillets were incubated with 1:100 LysoTracker Deep Red (ThermoFisher Scientific, cat L12492) in PBS with no detergent for 10 mins before fixation and mounting. Subsequent processing for IHC was conducted as indicated above with the exception that no detergent was used. Samples were mounted in Vectashield (Vector Labs) for spinning disk confocal imaging and in Prolong Diamond (ThermoFisher Scientific cat no. P36965) for SIM.

NMJs on muscle 6/7 from segments A2 and A3 were selected for EV and colocalization analyses, and muscle 4 from the same segments was used for bouton counting. For fluorescent intensity quantification, Spinning Disk Confocal stacks were collected at room temperature, using Nikon Elements AR software, on a Nikon Ni-E upright microscope equipped with 60X (n.a. 1.4) and 100X (n.a. 1.45) oil immersion objectives, a Yokogawa CSU-W1 spinning disk head, and an Andor iXon 897U EMCCD camera. Images for each independent experiment were acquired and are shown with identical settings for all conditions, except where indicated otherwise, and for all α-HRP immunostaining (which were used to analyze morphology and not for intensity quantifications).

3D-Structured Illumination Microscopy (SIM) was performed on a Nikon N-SIM E system (consisting of an inverted Eclipse Ti-E microscope, 100x (n.a 1.45) oil-immersion objective, and a Hamamatsu OrcaFLASH4 sCMOS camera). Channel alignment was calculated for each imaging session using tetraspeck beads (Invitrogen cat no. T-7284). Images were collected at room temperature with a regime of 3 grid orientations and 5 phases and were reconstructed using Nikon Elements software, using a theoretical, ideal OTF generated by the software.

### Image analysis & Statistics

3D analyses of presynaptic and postsynaptic EV cargo at NMJs were conducted using Volocity 6.0 software (Perkin Elmer, Waltham, MA). Manual thresholding of the α-HRP signal was used to define presynaptic volume (HRP-positive objects > 7 µm3, with holes filled) and postsynaptic volume (a dilation of 3 µm around the presynaptic volume).

The signal to be measured (e.g. APP-EGFP, Syt4-EGFP, or Rab7^KI^) was manually thresholded to distinguish postsynaptic puncta from background, and sum intensity measurements were made from APP-positive objects intersecting with either presynaptic or postsynaptic volumes as defined above. All intensity values were then normalized to the presynaptic volume. Puncta number and volume were measured in presynaptic volumes that were thresholded and defined as above.

Analysis of APP-EGFP in axons and motor neuron cell bodies was conducted using ImageJ. Axon intensity measurements were conducted from MaxIPs; HRP was used as a threshold to select a region of axon. APP intensity in motor neuron cell bodies was measured from a middle slice through each cell body.

Co-localization between APP and Rab11 was performed in Volocity. As above, the α-HRP channel was used as a mask to highlight the presynaptic area. For quantitative analysis of SIM images, the following unbiased criteria were applied to select images with sufficiently high-quality staining and signal:noise ratio, and that were adequately free of reconstruction artifacts. Both channels required a presynaptic CoV > 0.1, and ratio of presynaptic APP CoV:postsynaptic APP CoV of >1.5. A threshold was set to identify all APP or Rab11-positive volumes in the HRP area, and the fraction of the sum intensity of APP signal that coincided with Rab11 signal was calculated and vice versa.

### S2 cell culture and EV fractionation

S2 cells were cultured according to standard protocols (Cherbas and Cherbas, 2007) in Schneider’s media supplemented with 10% fetal bovine serum, 0.1 mg/mL penicillin/streptomycin (Lonza), and 30 µg/ml blasticidin. To create a stable cell line, pBI-UASc-APP^695^-EGFP was co-transfected using Effectene reagent (Qiagen) with pACSwitch (gift from M. Marr, based on (Marr et al., 2012), consisting of a 445bp version of the *Drosophila* Actin5C promoter driving the Geneswitch transcription factor (Roman et al., 2001) and a blasticidin resistance marker). 2 days after transfection, APP-EGFP expression was induced for 2 further days with 0.1 mm RU486, before cells were harvested.

To isolate EVs from cell culture supernatants, cells were harvested and centrifuged for 10 min at room temperature at 300 x *g* in a swinging bucket rotor. Pellets (representing total cellular lysate) were washed once in 1 ml PBS, then resuspended in 250 µl Laemmli sample buffer and boiled for 1 min. Culture supernatants were centrifuged for 10 min at 2000xg in a swinging bucket rotor, then for 30 min at 10000 x *g* in a fixed angle rotor. Supernatants were then transferred to ultracentrifuge tubes, supplemented with PBS to 32 ml, and centrifuged for 70 min at 100000 x *g* in a Beckman Ti70 rotor. Pellets were resuspended in 1 ml PBS, and centrifuged for 70 min at 100000 x *g* in a Beckman TLA100.3 rotor. 100000 x g pellets (containing EVs) were resuspended in 50 µl PBS, denatured with 50 µl Laemmli sample buffer, and boiled for 1 min. Samples were fractionated by SDS-PAGE, transferred to nitrocellulose, immunoblotted with the indicated primary antibodies and infrared dye conjugated secondary antibodies, and imaged on a LICOR Odyssey device.

To fractionate EVs on sucrose gradients, 100 ml of cell culture were centrifuged for 10 min at room temperature at 300 x *g* in a swinging bucket rotor. Culture supernatants were re-centrifuged for 10 min at 2000 x *g* in a swinging bucket rotor, then for 30 min at 10000 x *g* in a fixed angle rotor. Supernatants were then transferred to ultracentrifuge tubes, supplemented with PBS to 32 ml, and centrifuged for 70 min at 100000 x *g* in a Beckman Ti70 rotor. Pellets were resuspended in 1 ml PBS, and centrifuged for 70 min at 100000 x *g* in a Beckman TLA100.3 rotor. 100000 x *g* pellets were resuspended using a .22-gauge needle in 20 mM HEPES pH 7.5, 2.5 M sucrose, and a 10 ml discontinuous 2.5 – 0.25 M sucrose gradient in 20 mM HEPES pH 7.5 was layered above it. Gradients were centrifuged at 200,000 x *g* in a Sw40Ti rotor for 16 h. 1 ml fractions were collected from the bottom, and 20 µl of each fraction was weighed to determine density. 2 ml 20 mM HEPES was added to each fraction, and samples were centrifuged at 110,000 x *g* for 1 h. Pellets were resuspended in 20 µl PBS for immunoblotting and electron microscopy.

For electron microscopy of purified EVs, sucrose gradient fractions were diluted 1:5 into PBS, applied to copper grids coated with continuous carbon, negatively stained with 2% uranyl acetate (JT Baker Chemical Co., Phillipsburg, NJ), and imaged using an FEI Morgagni transmission electron microscope (FEI, Hillsboro, OR) operating at 80 kV and equipped with a 1k × 1k charge coupled device (CCD) camera (Gatan, Pleasanton, CA).

### Electron microscopy of larval NMJs

Wandering third-instar larvae (3 animals per condition) were dissected and fixed in 2.5% glutaraldehyde, 2% paraformaldehyde in 1% sodium cacodylate buffer overnight at 4ºC. Samples were postfixed in 1% Osmium Tetroxide, 1.5% Potassium ferrocyanide for 1h, then 1% aqueous Uranyl Acetate for 1 h. Stepwise dehydration was conducted for 10 minutes each in 50%, 70%, 90% ethanol. followed by 2x 10 minutes in 100% Ethanol. Samples were transferred to 100% Propylenoxide for 1 hour and then left overnight in a 1:1 mixture of Propylenoxide and 812 TAAB Epon Resin (TAAB Laboratories Equipment Ltd, Aldermaston, England). Samples were then transferred to fresh Epon, flat embedded and polymerized at 60ºC for 72 h, and remounted for sectioning. 70-µm thin sections were cut on a Leica UC6 Ultramicrotome (Leica Microsystems, Buffalo Grove, IL), collected on to copper grids coated with formvar and carbon and then poststained with Lead Citrate (Venable and Coggeshall, 1965). Grids were imaged using a FEI Morgagni TEM (see above) at 4400-5600x magnification, or on a CM12 transmission electron microscope operating at 120keV and equipped with LaB6 electron source and a Gatan Ultrascan 2kx2k CCD camera.

EM image analysis was performed on blinded images using FIJI. For quantification of endosomes and MVBs, endosomes were defined as presynaptic unilamellar compartments > 100 nm at its longest axis. MVBs were defined as endosomes that contained 1 or more 50-100 nm intraluminal vesicles. Extracellular vesicles were measured as objects that were between 50-100 nm in diameter, within 1 µm of the neuron, and apparently free-floating (no contiguity with the SSR membrane). Number of endosomes, MVBs, and EVs were normalized to the area of the bouton.

### Statistical methods

All statistical measurements were performed in GraphPad Prism 6 (see **Table 3**). Comparisons were made separately for presynaptic and postsynaptic datasets, due to differences between these compartments for intensity, signal-to-noise ratio and variance. Datasets were tested for normality, and statistical significance tested using unpaired Student’s t-tests or Mann-Whitney tests (if # conditions = 2) or ANOVA followed by Tukey’s tests or Kruskal-Wallis followed by Dunn’s test (if # conditions > 2).

## END MATTER

### Author contribution and notes

Author contributions: RBW, ANB, MJZ, and AAR designed the study and experiments. RBW, ANB, MJZ, ECD, SML, SW, BI, AY, and AAR conducted the experiments. RBW, ANB, MJZ, ECD, SML, SW, AY, and AAR performed the analyses. KK generated critical reagents. RBW and AAR wrote the manuscript. This article contains supporting information (3 supplemental Figures and 3 tables) online.

## Acknowledgements

We thank the Bloomington Drosophila Stock Center (NIH P40OD018537), the Developmental Studies Hybridoma Bank created by the NICHD of the NIH, Konrad Bassler, Vivian Budnik, Pete Cullen, Xinhua Lin, Kalpana White, and Philip Copenhaver for strains and reagents, and Crystal Yu, Troy Zhao, Agnieska Collins, and the Brandeis Electron Microscopy Facility for technical assistance. This work was supported by NINDS grants DP2 NS082127 and R01 NS103967 to A.A.R., T32 GM007122 to R.B.W., T32 NS007292 to E.C.D., and by the Brandeis NSF MRSEC, Bioinspired Soft Materials (NSF-DMR 2011846).

## REFERENCES

Akbergenova, Y., and J.T. Littleton. 2017. Pathogenic Huntington Alters BMP Signaling and Synaptic Growth through Local Disruptions of Endosomal Compartments. J Neurosci. 37:3425–3439.

Arbo, B.D., L.R. Cechinel, R.P. Palazzo, and I.R. Siqueira. 2020. Endosomal dysfunction impacts extracellular vesicle release: Central role in Abeta pathology. Ageing Res Rev. 58:101006.

Arighi, C.N., L.M. Hartnell, R.C. Aguilar, C.R. Haft, and J.S. Bonifacino. 2004. Role of the mammalian retromer in sorting of the cation-independent mannose 6-phosphate receptor. J Cell Biol. 165:123–133.

Ashley, J., B. Cordy, D. Lucia, L.G. Fradkin, V. Budnik, and T. Thomson. 2018. Retrovirus-like Gag Protein Arc1 Binds RNA and Traffics across Synaptic Boutons. Cell. 172:262–274 e211.

Baietti, M.F., Z. Zhang, E. Mortier, A. Melchior, G. Degeest, A. Geeraerts, Y. Ivarsson, F. Depoortere, C. Coomans, E. Vermeiren, P. Zimmermann, and G. David. 2012. Syndecan-syntenin-ALIX regulates the biogenesis of exosomes. Nat Cell Biol. 14:677–685.

Beckett, K., S. Monier, L. Palmer, C. Alexandre, H. Green, E. Bonneil, G. Raposo, P. Thibault, R. Le Borgne, and J.P. Vincent. 2013. *Drosophila* S2 cells secrete wingless on exosome-like vesicles but the wingless gradient forms independently of exosomes. Traffic. 14:82–96.

Becot, A., C. Volgers, and G. van Niel. 2020. Transmissible Endosomal Intoxication: A Balance between Exosomes and Lysosomes at the Basis of Intercellular Amyloid Propagation. Biomedicines. 8:272.

Berman, D.E., D. Ringe, G.A. Petsko, and S.A. Small. 2015. The use of pharmacological retromer chaperones in Alzheimer’s disease and other endosomal-related disorders. Neurotherapeutics. 12:12–18.

Bhalla, A., C.P. Vetanovetz, E. Morel, Z. Chamoun, G. Di Paolo, and S.A. Small. 2012. The location and trafficking routes of the neuronal retromer and its role in amyloid precursor protein transport. Neurobiol Dis. 47:126–134.

Brodin, L., and O. Shupliakov. 2018. Retromer in Synaptic Function and Pathology. Front Synaptic Neurosci. 10:37.

Buggia-Prevot, V., C.G. Fernandez, S. Riordan, K.S. Vetrivel, J. Roseman, J. Waters, V.P. Bindokas, R. Vassar, and G. Thinakaran. 2014. Axonal BACE1 dynamics and targeting in hippocampal neurons: a role for Rab11 GTPase. Mol Neurodegener. 9:1.

Cassar, M., and D. Kretzschmar. 2016. Analysis of Amyloid Precursor Protein Function in *Drosophila melanogaster*. Front Mol Neurosci. 9:61.

Chen, C., D. Garcia-Santos, Y. Ishikawa, A. Seguin, L. Li, K.H. Fegan, G.J. Hildick-Smith, D.I. Shah, J.D. Cooney, W. Chen, M.J. King, Y.Y. Yien, I.J. Schultz, H. Anderson, A.J. Dalton, M.L. Freedman, P.D. Kingsley, J. Palis, S.M. Hattangadi, H.F. Lodish, D.M. Ward, J. Kaplan, T. Maeda, P. Ponka, and B.H. Paw. 2013. Snx3 regulates recycling of the transferrin receptor and iron assimilation. Cell Metab. 17:343–352.

Chen, H.M., Y. Huang, B.D. Pfeiffer, X. Yao, and T. Lee. 2015. An enhanced gene targeting toolkit for *Drosophila*: Golic+. Genetics. 199:683–694.

Chen, K.E., M.D. Healy, and B.M. Collins. 2019a. Towards a molecular understanding of endosomal trafficking by Retromer and Retriever. Traffic. 20: 465–478.

Chen, X., J.K. Kordich, E.T. Williams, N. Levine, A. Cole-Strauss, L. Marshall, V. Labrie, J. Ma, J.W. Lipton, and D.J. Moore. 2019b. Parkinson’s disease-linked D620N VPS35 knockin mice manifest tau neuropathology and dopaminergic neurodegeneration. Proc Natl Acad Sci U S A. 116: 5765–5774.

Cherbas, L., and P. Cherbas. 2007. *Drosophila* cell culture and transformation. CSH Protoc. 2007:pdb top6.

Choy, R.W., Z. Cheng, and R. Schekman. 2012. Amyloid precursor protein (APP) traffics from the cell surface via endosomes for amyloid beta (Abeta) production in the trans-Golgi network. Proc Natl Acad Sci U S A. 109:E2077–82.

Choy, R.W., M. Park, P. Temkin, B.E. Herring, A. Marley, R.A. Nicoll, and M. von Zastrow. 2014. Retromer mediates a discrete route of local membrane delivery to dendrites. Neuron. 82:55–62.

Corrigan, L., S. Redhai, A. Leiblich, S.J. Fan, S.M. Perera, R. Patel, C. Gandy, S.M. Wainwright, J.F. Morris, F. Hamdy, D.C. Goberdhan, and C. Wilson. 2014. BMP-regulated exosomes from *Drosophila* male reproductive glands reprogram female behavior. J Cell Biol. 206:671–688.

Coulter, M.E., C.M. Dorobantu, G.A. Lodewijk, F. Delalande, S. Cianferani, V.S. Ganesh, R.S. Smith, E.T. Lim, C.S. Xu, S. Pang, E.T. Wong, H.G.W. Lidov, M.L. Calicchio, E. Yang, D.M. Gonzalez, T.M. Schlaeger, G.H. Mochida, H. Hess, W.A. Lee, M.K. Lehtinen, T. Kirchhausen, D. Haussler, F.M.J. Jacobs, R. Gaudin, and C.A. Walsh. 2018. The ESCRT-III Protein CHMP1A Mediates Secretion of Sonic Hedgehog on a Distinctive Subtype of Extracellular Vesicles. Cell Rep. 24:973–986 e978.

Cuartero, Y., M. Mellado, A. Capell, M. Alvarez-Dolado, and M. Verges. 2012. Retromer regulates postendocytic sorting of beta-secretase in polarized Madin-Darby canine kidney cells. Traffic. 13:1393–1410.

Cui, Y., J.M. Carosi, Z. Yang, N. Ariotti, M.C. Kerr, R.G. Parton, T.J. Sargeant, and R.D. Teasdale. 2019. Retromer has a selective function in cargo sorting via endosome transport carriers. J Cell Biol. 218:615–631.

Das, U., D.A. Scott, A. Ganguly, E.H. Koo, Y. Tang, and S. Roy. 2013. Activity-induced convergence of APP and BACE-1 in acidic microdomains via an endocytosis-dependent pathway. Neuron. 79:447–460.

Das, U., L. Wang, A. Ganguly, J.M. Saikia, S.L. Wagner, E.H. Koo, and S. Roy. 2016. Visualizing APP and BACE-1 approximation in neurons yields insight into the amyloidogenic pathway. Nat Neurosci. 19:55–64.

Derivery, E., E. Helfer, V. Henriot, and A. Gautreau. 2012. Actin polymerization controls the organization of WASH domains at the surface of endosomes. PLoS One. 7:e39774.

Dermaut, B., K.K. Norga, A. Kania, P. Verstreken, H. Pan, Y. Zhou, P. Callaerts, and H.J. Bellen. 2005. Aberrant lysosomal carbohydrate storage accompanies endocytic defects and neurodegeneration in *Drosophila* benchwarmer. J Cell Biol. 170:127–139.

Deshpande, M., Z. Feiger, A.K. Shilton, C.C. Luo, E. Silverman, and A.A. Rodal. 2016. Role of BMP receptor traffic in synaptic growth defects in an ALS model. Mol Biol Cell. 27:2898–2910.

Deshpande, M., and A.A. Rodal. 2016. The Crossroads of Synaptic Growth Signaling, Membrane Traffic and Neurological Disease: Insights from *Drosophila*. Traffic. 17:87–101.

Dolev, I., H. Fogel, H. Milshtein, Y. Berdichevsky, N. Lipstein, N. Brose, N. Gazit, and I. Slutsky. 2013. Spike bursts increase amyloid-beta 40/42 ratio by inducing a presenilin-1 conformational change. Nat Neurosci. 16:587–595.

Eggert, S., C. Thomas, S. Kins, and G. Hermey. 2018. Trafficking in Alzheimer’s Disease: Modulation of APP Transport and Processing by the Transmembrane Proteins LRP1, SorLA, SorCS1c, Sortilin, and Calsyntenin. Mol Neurobiol. 55:5809–5829.

Eitan, E., C. Suire, S. Zhang, and M.P. Mattson. 2016. Impact of lysosome status on extracellular vesicle content and release. Ageing Res Rev. 32:65–74.

Escudero, C.A., O.M. Lazo, C. Galleguillos, J.I. Parraguez, M.A. Lopez-Verrilli, C. Cabeza, L. Leon, U. Saeed, C. Retamal, A. Gonzalez, M.P. Marzolo, B.D. Carter, F.A. Court, and F.C. Bronfman. 2014. The p75 neurotrophin receptor evades the endolysosomal route in neuronal cells, favouring multivesicular bodies specialised for exosomal release. J Cell Sci. 127:1966–1979.

Fan, S.-J., B. Kroeger, P.P. Marie, E.M. Bridges, J.D. Mason, K. McCormick, C. Zois, H. Sheldon, N.K. Alham, E. Johnson, M. Ellis, M.I. Stefana, C.C. Mendes, S.M. Wainwright, C. Cunningham, F.C. Hamdy, J.F. Morris, A.L. Harris, C. Wilson, and D.C.I. Goberdhan. 2019. Glutamine Deprivation Regulates the Origin and Function of Cancer Cell Exosomes. bioRxiv. https://doi.org/10.1101/859447

Faure, J., G. Lachenal, M. Court, J. Hirrlinger, C. Chatellard-Causse, B. Blot, J. Grange, G. Schoehn, Y. Goldberg, V. Boyer, F. Kirchhoff, G. Raposo, J. Garin, and R. Sadoul. 2006. Exosomes are released by cultured cortical neurones. Mol Cell Neurosci. 31:642–648.

Feng, Y., A. Ueda, and C.F. Wu. 2004. A modified minimal hemolymph-like solution, HL3.1, for physiological recordings at the neuromuscular junctions of normal and mutant *Drosophila* larvae. J Neurogenet. 18:377–402.

Ferguson, S.M. 2018. Axonal transport and maturation of lysosomes. Curr Opin Neurobiol. 51:45–51.

Fernandes, A.C., V. Uytterhoeven, S. Kuenen, Y.C. Wang, J.R. Slabbaert, J. Swerts, J. Kasprowicz, S. Aerts, and P. Verstreken. 2014. Reduced synaptic vesicle protein degradation at lysosomes curbs TBC1D24/sky-induced neurodegeneration. J Cell Biol. 207:453–462.

Fjorback, A.W., M. Seaman, C. Gustafsen, A. Mehmedbasic, S. Gokool, C. Wu, D. Militz, V. Schmidt, P. Madsen, J.R. Nyengaard, T.E. Willnow, E.I. Christensen, W.B. Mobley, A. Nykjaer, and O.M. Andersen. 2012. Retromer binds the FANSHY sorting motif in SorLA to regulate amyloid precursor protein sorting and processing. J Neurosci. 32:1467–1480.

Follett, J., S.J. Norwood, N.A. Hamilton, M. Mohan, O. Kovtun, S. Tay, Y. Zhe, S.A. Wood, G.D. Mellick, P.A. Silburn, B.M. Collins, A. Bugarcic, and R.D. Teasdale. 2014. The Vps35 D620N mutation linked to Parkinson’s disease disrupts the cargo sorting function of retromer. Traffic. 15:230–244.

French, K.C., M.A. Antonyak, and R.A. Cerione. 2017. Extracellular vesicle docking at the cellular port: Extracellular vesicle binding and uptake. Semin Cell Dev Biol. 67:48–55.

Fuentes-Medel, Y., M.A. Logan, J. Ashley, B. Ataman, V. Budnik, and M.R. Freeman. 2009. Glia and muscle sculpt neuromuscular arbors by engulfing destabilized synaptic boutons and shed presynaptic debris. PLoS Biol. 7:e1000184.

Gong, J., R. Korner, L. Gaitanos, and R. Klein. 2016. Exosomes mediate cell contact-independent ephrin-Eph signaling during axon guidance. J Cell Biol. 214:35–44.

Gonzalez-Noriega, A., J.H. Grubb, V. Talkad, and W.S. Sly. 1980. Chloroquine inhibits lysosomal enzyme pinocytosis and enhances lysosomal enzyme secretion by impairing receptor recycling. J Cell Biol. 85:839–852.

Gross, J.C., V. Chaudhary, K. Bartscherer, and M. Boutros. 2012. Active Wnt proteins are secreted on exosomes. Nat Cell Biol. 14:1036–1045.

Gunawardena, S., and L.S. Goldstein. 2001. Disruption of axonal transport and neuronal viability by amyloid precursor protein mutations in *Drosophila*. Neuron. 32:389–401.

Harbour, M.E., S.Y. Breusegem, R. Antrobus, C. Freeman, E. Reid, and M.N. Seaman. 2010. The cargo-selective retromer complex is a recruiting hub for protein complexes that regulate endosomal tubule dynamics. J Cell Sci. 123:3703–3717.

Holm, M.M., J. Kaiser, and M.E. Schwab. 2018. Extracellular Vesicles: Multimodal Envoys in Neural Maintenance and Repair. Trends Neurosci. 41:360–372.

Hussain, N.K., G.H. Diering, J. Sole, V. Anggono, and R.L. Huganir. 2014. Sorting Nexin 27 regulates basal and activity-dependent trafficking of AMPARs. Proc Natl Acad Sci U S A. 111:11840–11845.

Inoshita, T., T. Arano, Y. Hosaka, H. Meng, Y. Umezaki, S. Kosugi, T. Morimoto, M. Koike, H.Y. Chang, Y. Imai, and N. Hattori. 2017. Vps35 in cooperation with LRRK2 regulates synaptic vesicle endocytosis through the endosomal pathway in *Drosophila*. Hum Mol Genet. 26: 2933–2948.

Jia, D., T.S. Gomez, D.D. Billadeau, and M.K. Rosen. 2012. Multiple repeat elements within the FAM21 tail link the WASH actin regulatory complex to the retromer. Mol Biol Cell. 23:2352–2361.

Jimenez-Orgaz, A., A. Kvainickas, H. Nagele, J. Denner, S. Eimer, J. Dengjel, and F. Steinberg. 2017. Control of RAB7 activity and localization through the retromer-TBC1D5 complex enables RAB7-dependent mitophagy. EMBO J. 37:235–254.

Kanmert, D., A. Cantlon, C.R. Muratore, M. Jin, T.T. O’Malley, G. Lee, T.L. Young-Pearse, D.J. Selkoe, and D.M. Walsh. 2015. C-Terminally Truncated Forms of Tau, But Not Full-Length Tau or Its C-Terminal Fragments, Are Released from Neurons Independently of Cell Death. J Neurosci. 35:10851–10865.

Koles, K., J. Nunnari, C. Korkut, R. Barria, C. Brewer, Y. Li, J. Leszyk, B. Zhang, and V. Budnik. 2012. Mechanism of evenness interrupted (Evi)-exosome release at synaptic boutons. J Biol Chem. 287:16820–16834.

Koles, K., A.R. Yeh, and A.A. Rodal. 2015. Tissue-specific tagging of endogenous loci in *Drosophila melanogaster*. Biol Open. 5:83–89.

Korkut, C., B. Ataman, P. Ramachandran, J. Ashley, R. Barria, N. Gherbesi, and V. Budnik. 2009. Trans-synaptic transmission of vesicular Wnt signals through Evi/Wntless. Cell. 139:393–404.

Korkut, C., Y. Li, K. Koles, C. Brewer, J. Ashley, M. Yoshihara, and V. Budnik. 2013. Regulation of postsynaptic retrograde signaling by presynaptic exosome release. Neuron. 77:1039–1046.

Korolchuk, V.I., M.M. Schutz, C. Gomez-Llorente, J. Rocha, N.R. Lansu, S.M. Collins, Y.P. Wairkar, I.M. Robinson, and C.J. O’Kane. 2007. *Drosophila* Vps35 function is necessary for normal endocytic trafficking and actin cytoskeleton organisation. J Cell Sci. 120:4367–4376.

Kvainickas, A., H. Nagele, W. Qi, L. Dokladal, A. Jimenez-Orgaz, L. Stehl, D. Gangurde, Q. Zhao, Z. Hu, J. Dengjel, C. De Virgilio, R. Baumeister, and F. Steinberg. 2019. Retromer and TBC1D5 maintain late endosomal RAB7 domains to enable amino acid-induced mTORC1 signaling. J Cell Biol. 218:3019–3038.

Laulagnier, K., C. Javalet, F.J. Hemming, M. Chivet, G. Lachenal, B. Blot, C. Chatellard, and R. Sadoul. 2017. Amyloid precursor protein products concentrate in a subset of exosomes specifically endocytosed by neurons. Cell Mol Life Sci. 75:757–773.

Lauwers, E., Y.C. Wang, R. Gallardo, R. Van der Kant, E. Michiels, J. Swerts, P. Baatsen, S.S. Zaiter, S.R. McAlpine, N.V. Gounko, F. Rousseau, J. Schymkowitz, and P. Verstreken. 2018. Hsp90 Mediates Membrane Deformation and Exosome Release. Mol Cell. 71:689–702 e689.

Lazarov, O., M. Lee, D.A. Peterson, and S.S. Sisodia. 2002. Evidence that synaptically released beta-amyloid accumulates as extracellular deposits in the hippocampus of transgenic mice. J Neurosci. 22:9785–9793.

Li, J., T. Kanekiyo, M. Shinohara, Y. Zhang, M.J. LaDu, H. Xu, and G. Bu. 2012. Differential regulation of amyloid-beta endocytic trafficking and lysosomal degradation by apolipoprotein E isoforms. J Biol Chem. 287:44593–44601.

Li, J.G., J. Chiu, and D. Pratico. 2019. Full recovery of the Alzheimer’s disease phenotype by gain of function of vacuolar protein sorting 35. Mol Psychiatry. 25: 2630–2640.

Loo, L.S., N. Tang, M. Al-Haddawi, G.S. Dawe, and W. Hong. 2014. A role for sorting nexin 27 in AMPA receptor trafficking. Nat Commun. 5:3176.

Lundgren, J.L., S. Ahmed, B. Winblad, G.K. Gouras, L.O. Tjernberg, and S. Frykman. 2014. Activity-independent release of the amyloid beta-peptide from rat brain nerve terminals. Neurosci Lett. 566:125–130.

Luo, L.Q., L.E. Martin-Morris, and K. White. 1990. Identification, secretion, and neural expression of APPL, a *Drosophila* protein similar to human amyloid protein precursor. J Neurosci. 10:3849–3861.

Malik, B.R., V.K. Godena, and A.J. Whitworth. 2015. VPS35 pathogenic mutations confer no dominant toxicity but partial loss of function in *Drosophila* and genetically interact with parkin. Hum Mol Genet. 24:6106–6117.

Marie, P.P., S.-J. Fan, C.C. Mendes, S.M. Wainwright, A.L. Harris, D.C.I. Goberdhan, and C. Wilson. 2020. Accessory ESCRT-III proteins selectively regulate Rab11-exosome biogenesis in *Drosophila* secondary cells. bioRxiv. doi: https://doi.org/10.1101/2020.06.18.158725

Marr, S.K., K.L. Pennington, and M.T. Marr. 2012. Efficient metal-specific transcription activation by *Drosophila* MTF-1 requires conserved cysteine residues in the carboxy-terminal domain. Biochim Biophys Acta. 1819:902–912.

Maruzs, T., P. Lorincz, Z. Szatmari, S. Szeplaki, Z. Sandor, Z. Lakatos, G. Puska, G. Juhasz, and M. Sass. 2015. Retromer Ensures the Degradation of Autophagic Cargo by Maintaining Lysosome Function in *Drosophila*. Traffic. 16:1088–1107.

McGough, I.J., F. Steinberg, D. Jia, P.A. Barbuti, K.J. McMillan, K.J. Heesom, A.L. Whone, M.A. Caldwell, D.D. Billadeau, M.K. Rosen, and P.J. Cullen. 2014. Retromer binding to FAM21 and the WASH complex is perturbed by the Parkinson disease-linked VPS35(D620N) mutation. Curr Biol. 24:1670–1676.

McNally, K.E., and P.J. Cullen. 2018. Endosomal Retrieval of Cargo: Retromer Is Not Alone. Trends Cell Biol. 28:807–822.

Mecozzi, V.J., D.E. Berman, S. Simoes, C. Vetanovetz, M.R. Awal, V.M. Patel, R.T. Schneider, G.A. Petsko, D. Ringe, and S.A. Small. 2014. Pharmacological chaperones stabilize retromer to limit APP processing. Nat Chem Biol. 10:443–449.

Meresse, S., J.P. Gorvel, and P. Chavrier. 1995. The rab7 GTPase resides on a vesicular compartment connected to lysosomes. J Cell Sci. 108 (Pt 11):3349–3358.

Miranda, A.M., Z.M. Lasiecka, Y. Xu, J. Neufeld, S. Shahriar, S. Simoes, R.B. Chan, T.G. Oliveira, S.A. Small, and G. Di Paolo. 2018. Neuronal lysosomal dysfunction releases exosomes harboring APP C-terminal fragments and unique lipid signatures. Nat Commun. 9:291.

Miura, E., T. Hasegawa, M. Konno, M. Suzuki, N. Sugeno, N. Fujikake, S. Geisler, M. Tabuchi, R. Oshima, A. Kikuchi, T. Baba, K. Wada, Y. Nagai, A. Takeda, and M. Aoki. 2014. VPS35 dysfunction impairs lysosomal degradation of alpha-synuclein and exacerbates neurotoxicity in a *Drosophila* model of Parkinson’s disease. Neurobiol Dis. 71:1–13.

Muhammad, A., I. Flores, H. Zhang, R. Yu, A. Staniszewski, E. Planel, M. Herman, L. Ho, R. Kreber, L.S. Honig, B. Ganetzky, K. Duff, O. Arancio, and S.A. Small. 2008. Retromer deficiency observed in Alzheimer’s disease causes hippocampal dysfunction, neurodegeneration, and Abeta accumulation. Proc Natl Acad Sci U S A. 105:7327–7332.

Munsie, L.N., A.J. Milnerwood, P. Seibler, D.A. Beccano-Kelly, I. Tatarnikov, J. Khinda, M. Volta, C. Kadgien, L.P. Cao, L. Tapia, C. Klein, and M.J. Farrer. 2015. Retromer-dependent neurotransmitter receptor trafficking to synapses is altered by the Parkinson’s disease VPS35 mutation p.D620N. Hum Mol Genet. 24:1691–1703.

Nakano, Y. 2019. Stories of spinster with various faces: from courtship rejection to tumor metastasis rejection. J Neurogenet:1–6.

Narayanan, R., H. Kramer, and M. Ramaswami. 2000. *Drosophila* endosomal proteins hook and deep orange regulate synapse size but not synaptic vesicle recycling. J Neurobiol. 45:105–119.

Naslavsky, N., and S. Caplan. 2018. The enigmatic endosome - sorting the ins and outs of endocytic trafficking. J Cell Sci. 131.

Neuman, S.D., E.L. Terry, J.E. Selegue, A.T. Cavanagh, and A. Bashirullah. 2020. Mistargeting of secretory cargo in retromer-deficient cells. bioRxiv. doi: https://doi.org/10.1101/2020.07.02.185660

Ni, J.Q., M. Markstein, R. Binari, B. Pfeiffer, L.P. Liu, C. Villalta, M. Booker, L. Perkins, and N. Perrimon. 2008. Vector and parameters for targeted transgenic RNA interference in *Drosophila melanogaster*. Nature methods. 5:49–51.

Nielsen, M.S., C. Gustafsen, P. Madsen, J.R. Nyengaard, G. Hermey, O. Bakke, M. Mari, P. Schu, R. Pohlmann, A. Dennes, and C.M. Petersen. 2007. Sorting by the cytoplasmic domain of the amyloid precursor protein binding receptor SorLA. Mol Cell Biol. 27:6842–6851.

Norris, A., P. Tammineni, S. Wang, J. Gerdes, A. Murr, K.Y. Kwan, Q. Cai, and B.D. Grant. 2017. SNX-1 and RME-8 oppose the assembly of HGRS-1/ESCRT-0 degradative microdomains on endosomes. Proc Natl Acad Sci U S A. 114:E307–E316.

O’Connor-Giles, K.M., L.L. Ho, and B. Ganetzky. 2008. Nervous wreck interacts with thickveins and the endocytic machinery to attenuate retrograde BMP signaling during synaptic growth. Neuron. 58:507–518.

Pastuzyn, E.D., C.E. Day, R.B. Kearns, M. Kyrke-Smith, A.V. Taibi, J. McCormick, N. Yoder, D.M. Belnap, S. Erlendsson, D.R. Morado, J.A.G. Briggs, C. Feschotte, and J.D. Shepherd. 2018. The Neuronal Gene Arc Encodes a Repurposed Retrotransposon Gag Protein that Mediates Intercellular RNA Transfer. Cell. 172:275–288 e218.

Perez-Gonzalez, R., S.A. Gauthier, A. Kumar, and E. Levy. 2012. The exosome secretory pathway transports amyloid precursor protein carboxyl-terminal fragments from the cell into the brain extracellular space. J Biol Chem. 287:43108–43115.

Pocha, S.M., T. Wassmer, C. Niehage, B. Hoflack, and E. Knust. 2011. Retromer controls epithelial cell polarity by trafficking the apical determinant Crumbs. Curr Biol. 21:1111–1117.

Rajendran, L., M. Honsho, T.R. Zahn, P. Keller, K.D. Geiger, P. Verkade, and K. Simons. 2006. Alzheimer’s disease beta-amyloid peptides are released in association with exosomes. Proc Natl Acad Sci U S A. 103:11172–11177.

Ramaker, J.M., R.S. Cargill, T.L. Swanson, H. Quirindongo, M. Cassar, D. Kretzschmar, and P.F. Copenhaver. 2016. Amyloid Precursor Proteins Are Dynamically Trafficked and Processed during Neuronal Development. Front Mol Neurosci. 9:130.

Rink, J., E. Ghigo, Y. Kalaidzidis, and M. Zerial. 2005. Rab conversion as a mechanism of progression from early to late endosomes. Cell. 122:735–749.

Rodal, A.A., A.D. Blunk, Y. Akbergenova, R.A. Jorquera, L.K. Buhl, and J.T. Littleton. 2011. A presynaptic endosomal trafficking pathway controls synaptic growth signaling. J Cell Biol. 193:201–217.

Rojas, R., T. van Vlijmen, G.A. Mardones, Y. Prabhu, A.L. Rojas, S. Mohammed, A.J. Heck, G. Raposo, P. van der Sluijs, and J.S. Bonifacino. 2008. Regulation of retromer recruitment to endosomes by sequential action of Rab5 and Rab7. J Cell Biol. 183:513–526.

Roman, G., K. Endo, L. Zong, and R.L. Davis. 2001. P[Switch], a system for spatial and temporal control of gene expression in *Drosophila* melanogaster. Proc Natl Acad Sci U S A. 98:12602–12607.

Sardar Sinha, M., A. Ansell-Schultz, L. Civitelli, C. Hildesjo, M. Larsson, L. Lannfelt, M. Ingelsson, and M. Hallbeck. 2018. Alzheimer’s disease pathology propagation by exosomes containing toxic amyloid-beta oligomers. Acta Neuropathol. 136:41–56.

Satoh, A.K., J.E. O’Tousa, K. Ozaki, and D.F. Ready. 2005. Rab11 mediates post-Golgi trafficking of rhodopsin to the photosensitive apical membrane of *Drosophila* photoreceptors. Development. 132:1487–1497.

Savina, A., M. Vidal, and M.I. Colombo. 2002. The exosome pathway in K562 cells is regulated by Rab11. J Cell Sci. 115:2505–2515.

Schedin-Weiss, S., I. Caesar, B. Winblad, H. Blom, and L.O. Tjernberg. 2016. Super-resolution microscopy reveals gamma-secretase at both sides of the neuronal synapse. Acta Neuropathol Commun. 4:29.

Seaman, M.N., A. Gautreau, and D.D. Billadeau. 2013. Retromer-mediated endosomal protein sorting: all WASHed up! Trends Cell Biol. 23:522–528.

Seaman, M.N., M.E. Harbour, D. Tattersall, E. Read, and N. Bright. 2009. Membrane recruitment of the cargo-selective retromer subcomplex is catalysed by the small GTPase Rab7 and inhibited by the Rab-GAP TBC1D5. J Cell Sci. 122:2371–2382.

Seaman, M.N., J.M. McCaffery, and S.D. Emr. 1998. A membrane coat complex essential for endosome-to-Golgi retrograde transport in yeast. J Cell Biol. 142:665–681.

Seaman, M.N.J., A.S. Mukadam, and S.Y. Breusegem. 2018. Inhibition of TBC1D5 activates Rab7a and can enhance the function of the retromer cargo-selective complex. J Cell Sci. 131. jcs217398.

Sharples, R.A., L.J. Vella, R.M. Nisbet, R. Naylor, K. Perez, K.J. Barnham, C.L. Masters, and A.F. Hill. 2008. Inhibition of gamma-secretase causes increased secretion of amyloid precursor protein C-terminal fragments in association with exosomes. FASEB J. 22:1469–1478.

Simonetti, B., C.M. Danson, K.J. Heesom, and P.J. Cullen. 2017. Sequence-dependent cargo recognition by SNX-BARs mediates retromer-independent transport of CI-MPR. J Cell Biol. 216: 3695–3712.

Simonetti, B., B. Paul, K. Chaudhari, S. Weeratunga, F. Steinberg, M. Gorla, K.J. Heesom, G.J. Bashaw, B.M. Collins, and P.J. Cullen. 2019. Molecular identification of a BAR domain-containing coat complex for endosomal recycling of transmembrane proteins. Nat Cell Biol. 21:1219–1233.

Small, S.A., K. Kent, A. Pierce, C. Leung, M.S. Kang, H. Okada, L. Honig, J.P. Vonsattel, and T.W. Kim. 2005. Model-guided microarray implicates the retromer complex in Alzheimer’s disease. Ann Neurol. 58:909–919.

Small, S.A., S. Simoes-Spassov, R. Mayeux, and G.A. Petsko. 2017. Endosomal Traffic Jams Represent a Pathogenic Hub and Therapeutic Target in Alzheimer’s Disease. Trends Neurosci. 40:592–602.

Snow, P.M., N.H. Patel, A.L. Harrelson, and C.S. Goodman. 1987. Neural-specific carbohydrate moiety shared by many surface glycoproteins in *Drosophila* and grasshopper embryos. J Neurosci. 7:4137–4144.

Song, P., K. Trajkovic, T. Tsunemi, and D. Krainc. 2016. Parkin Modulates Endosomal Organization and Function of the Endo-Lysosomal Pathway. J Neurosci. 36:2425–2437.

Song, Z., Y. Xu, W. Deng, L. Zhang, H. Zhu, P. Yu, Y. Qu, W. Zhao, Y. Han, and C. Qin. 2020. Brain Derived Exosomes Are a Double-Edged Sword in Alzheimer’s Disease. Front Mol Neurosci. 13:79.

Steinberg, F., M. Gallon, M. Winfield, E.C. Thomas, A.J. Bell, K.J. Heesom, J.M. Tavare, and P.J. Cullen. 2013. A global analysis of SNX27-retromer assembly and cargo specificity reveals a function in glucose and metal ion transport. Nat Cell Biol. 15:461–471.

Stenmark, H., R.G. Parton, O. Steele-Mortimer, A. Lutcke, J. Gruenberg, and M. Zerial. 1994. Inhibition of rab5 GTPase activity stimulates membrane fusion in endocytosis. EMBO J. 13:1287–1296.

Strutt, H., P.F. Langton, N. Pearson, K.J. McMillan, D. Strutt, and P.J. Cullen. 2019. Retromer Controls Planar Polarity Protein Levels and Asymmetric Localization at Intercellular Junctions. Curr Biol. 29:484–491 e486.

Sullivan, C.P., A.G. Jay, E.C. Stack, M. Pakaluk, E. Wadlinger, R.E. Fine, J.M. Wells, and P.J. Morin. 2011. Retromer disruption promotes amyloidogenic APP processing. Neurobiol Dis. 43:338–345.

Swanson, T.L., L.M. Knittel, T.M. Coate, S.M. Farley, M.A. Snyder, and P.F. Copenhaver. 2005. The insect homologue of the amyloid precursor protein interacts with the heterotrimeric G protein Go alpha in an identified population of migratory neurons. Dev Biol. 288:160–178.

Sweeney, S.T., and G.W. Davis. 2002. Unrestricted synaptic growth in spinster-a late endosomal protein implicated in TGF-beta-mediated synaptic growth regulation. Neuron. 36:403–416.

Tan, J.Z.A., and P.A. Gleeson. 2019. The role of membrane trafficking in the processing of amyloid precursor protein and production of amyloid peptides in Alzheimer’s disease. Biochim Biophys Acta Biomembr. 1861:697–712.

Tang, F.L., W. Liu, J.X. Hu, J.R. Erion, J. Ye, L. Mei, and W.C. Xiong. 2015. VPS35 Deficiency or Mutation Causes Dopaminergic Neuronal Loss by Impairing Mitochondrial Fusion and Function. Cell Rep. 12:1631–1643.

Temkin, P., B. Lauffer, S. Jager, P. Cimermancic, N.J. Krogan, and M. von Zastrow. 2011. SNX27 mediates retromer tubule entry and endosome-to-plasma membrane trafficking of signalling receptors. Nat Cell Biol. 13:715–721.

Temkin, P., W. Morishita, D. Goswami, K. Arendt, L. Chen, and R. Malenka. 2017. The Retromer Supports AMPA Receptor Trafficking During LTP. Neuron. 94:74–82 e75.

Thery, C., K.W. Witwer, E. Aikawa, M.J. Alcaraz, J.D. Anderson, R. Andriantsitohaina, A. Antoniou, T. Arab, F. Archer, G.K. Atkin-Smith, D.C. Ayre, J.M. Bach, D. Bachurski, H. Baharvand, L. Balaj, S. Baldacchino, N.N. Bauer, A.A. Baxter, M. Bebawy, C. Beckham, A. Bedina Zavec, A. Benmoussa, A.C. Berardi, P. Bergese, E. Bielska, C. Blenkiron, S. Bobis-Wozowicz, E. Boilard, W. Boireau, A. Bongiovanni, F.E. Borras, S. Bosch, C.M. Boulanger, X. Breakefield, A.M. Breglio, M.A. Brennan, D.R. Brigstock, A. Brisson, M.L. Broekman, J.F. Bromberg, P. Bryl-Gorecka, S. Buch, A.H. Buck, D. Burger, S. Busatto, D. Buschmann, B. Bussolati, E.I. Buzas, J.B. Byrd, G. Camussi, D.R. Carter, S. Caruso, L.W. Chamley, Y.T. Chang, C. Chen, S. Chen, L. Cheng, A.R. Chin, A. Clayton, S.P. Clerici, A. Cocks, E. Cocucci, R.J. Coffey, A. Cordeiro-da-Silva, Y. Couch, F.A. Coumans, B. Coyle, R. Crescitelli, M.F. Criado, C. D’Souza-Schorey, S. Das, A. Datta Chaudhuri, P. de Candia, E.F. De Santana, O. De Wever, H.A. Del Portillo, T. Demaret, S. Deville, A. Devitt, B. Dhondt, D. Di Vizio, L.C. Dieterich, V. Dolo, A.P. Dominguez Rubio, M. Dominici, M.R. Dourado, T.A. Driedonks, F.V. Duarte, H.M. Duncan, R.M. Eichenberger, K. Ekstrom, S. El Andaloussi, C. Elie-Caille, U. Erdbrugger, J.M. Falcon-Perez, F. Fatima, J.E. Fish, M. Flores-Bellver, A. Forsonits, A. Frelet-Barrand, et al. 2018. Minimal information for studies of extracellular vesicles 2018 (MISEV2018): a position statement of the International Society for Extracellular Vesicles and update of the MISEV2014 guidelines. J Extracell Vesicles. 7:1535750.

Tian, Y., F.L. Tang, X. Sun, L. Wen, L. Mei, B.S. Tang, and W.C. Xiong. 2015. VPS35-deficiency results in an impaired AMPA receptor trafficking and decreased dendritic spine maturation. Mol Brain. 8:70.

Toh, W.H., J.Z. Tan, K.L. Zulkefli, F.J. Houghton, and P.A. Gleeson. 2017. Amyloid precursor protein traffics from the Golgi directly to early endosomes in an Arl5b- and AP4-dependent pathway. Traffic. 18:159–175.

Torroja, L., M. Packard, M. Gorczyca, K. White, and V. Budnik. 1999. The *Drosophila* beta-amyloid precursor protein homolog promotes synapse differentiation at the neuromuscular junction. J Neurosci. 19:7793–7803.

Udayar, V., V. Buggia-Prevot, R.L. Guerreiro, G. Siegel, N. Rambabu, A.L. Soohoo, M. Ponnusamy, B. Siegenthaler, J. Bali, Aesg, M. Simons, J. Ries, M.A. Puthenveedu, J. Hardy, G. Thinakaran, and L. Rajendran. 2013. A paired RNAi and RabGAP overexpression screen identifies Rab11 as a regulator of beta-amyloid production. Cell Rep. 5:1536–1551.

Ullrich, O., S. Reinsch, S. Urbe, M. Zerial, and R.G. Parton. 1996. Rab11 regulates recycling through the pericentriolar recycling endosome. J Cell Biol. 135:913–924.

van Niel, G., G. D’Angelo, and G. Raposo. 2018. Shedding light on the cell biology of extracellular vesicles. Nat Rev Mol Cell Biol. 19: 213–228.

Vardarajan, B.N., S.Y. Bruesegem, M.E. Harbour, R. Inzelberg, R. Friedland, P. St George-Hyslop, M.N. Seaman, and L.A. Farrer. 2012. Identification of Alzheimer disease-associated variants in genes that regulate retromer function. Neurobiol Aging. 33:2231 e2215–2231 e2230.

Vazquez-Sanchez, S., S. Bobeldijk, M.P. Dekker, L. van Keimpema, and J.R.T. van Weering. 2018. VPS35 depletion does not impair presynaptic structure and function. Sci Rep. 8:2996.

Venable, J.H., and R. Coggeshall. 1965. A Simplified Lead Citrate Stain for Use in Electron Microscopy. J Cell Biol. 25:407–408.

Vieira, S.I., S. Rebelo, H. Esselmann, J. Wiltfang, J. Lah, R. Lane, S.A. Small, S. Gandy, E.S.E.F. da Cruz, and E.S.O.A. da Cruz. 2010. Retrieval of the Alzheimer’s amyloid precursor protein from the endosome to the TGN is S655 phosphorylation state-dependent and retromer-mediated. Mol Neurodegener. 5:40.

Vingtdeux, V., M. Hamdane, A. Loyens, P. Gele, H. Drobeck, S. Begard, M.C. Galas, A. Delacourte, J.C. Beauvillain, L. Buee, and N. Sergeant. 2007. Alkalizing drugs induce accumulation of amyloid precursor protein by-products in luminal vesicles of multivesicular bodies. J Biol Chem. 282:18197–18205.

Wang, C.L., F.L. Tang, Y. Peng, C.Y. Shen, L. Mei, and W.C. Xiong. 2012a. VPS 35 regulates developing mouse hippocampal neuronal morphogenesis by promoting retrograde trafficking of BACE1. Biol Open. 1:1248–1257.

Wang, J., A. Fedoseienko, B. Chen, E. Burstein, D. Jia, and D.D. Billadeau. 2018. Endosomal receptor trafficking: Retromer and beyond. Traffic. 19:578–590.

Wang, J.W., E.S. Beck, and B.D. McCabe. 2012b. A modular toolset for recombination transgenesis and neurogenetic analysis of Drosophila. PLoS One. 7:e42102.

Wang, S., K.L. Tan, M.A. Agosto, B. Xiong, S. Yamamoto, H. Sandoval, M. Jaiswal, V. Bayat, K. Zhang, W.L. Charng, G. David, L. Duraine, K. Venkatachalam, T.G. Wensel, and H.J. Bellen. 2014. The retromer complex is required for rhodopsin recycling and its loss leads to photoreceptor degeneration. PLoS Biol. 12:e1001847.

Wang, X., Y. Zhao, X. Zhang, H. Badie, Y. Zhou, Y. Mu, L.S. Loo, L. Cai, R.C. Thompson, B. Yang, Y. Chen, P.F. Johnson, C. Wu, G. Bu, W.C. Mobley, D. Zhang, F.H. Gage, B. Ranscht, Y.W. Zhang, S.A. Lipton, W. Hong, and H. Xu. 2013. Loss of sorting nexin 27 contributes to excitatory synaptic dysfunction by modulating glutamate receptor recycling in Down’s syndrome. Nat Med. 19:473–480.

Wen, L., F.L. Tang, Y. Hong, S.W. Luo, C.L. Wang, W. He, C. Shen, J.U. Jung, F. Xiong, D.H. Lee, Q.G. Zhang, D. Brann, T.W. Kim, R. Yan, L. Mei, and W.C. Xiong. 2011. VPS35 haploinsufficiency increases Alzheimer’s disease neuropathology. J Cell Biol. 195:765–779.

Wilcke, M., L. Johannes, T. Galli, V. Mayau, B. Goud, and J. Salamero. 2000. Rab11 regulates the compartmentalization of early endosomes required for efficient transport from early endosomes to the trans-golgi network. J Cell Biol. 151:1207–1220.

Winckler, B., V. Faundez, S. Maday, Q. Cai, C. Guimas Almeida, and H. Zhang. 2018. The Endolysosomal System and Proteostasis: From Development to Degeneration. J Neurosci. 38:9364–9374.

Wu, S., R.R. Fagan, C. Uttamapinant, L.M. Lifshitz, K.E. Fogarty, A.Y. Ting, and H.E. Melikian. 2017. The Dopamine Transporter Recycles via a Retromer-Dependent Postendocytic Mechanism: Tracking Studies Using a Novel Fluorophore-Coupling Approach. J Neurosci. 37:9438–9452.

Ye, H., S.A. Ojelade, D. Li-Kroeger, Z. Zuo, L. Wang, Y. Li, J.Y. Gu, U. Tepass, A.A. Rodal, H.J. Bellen, and J.M. Shulman. 2020. Retromer subunit, VPS29, regulates synaptic transmission and is required for endolysosomal function in the aging brain. Elife. 9. e51977

Yong, X., L. Zhao, W. Deng, H. Sun, X. Zhou, L. Mao, W. Hu, X. Shen, Q. Sun, D.D. Billadeau, Y. Xue, and D. Jia. 2020. Mechanism of cargo recognition by retromer-linked SNX-BAR proteins. PLoS Biol. 18:e3000631.

Yu, Y., D.C. Jans, B. Winblad, L.O. Tjernberg, and S. Schedin-Weiss. 2018. Neuronal Abeta42 is enriched in small vesicles at the presynaptic side of synapses. Life Sci Alliance. 1:e201800028.

Zavodszky, E., M.N. Seaman, K. Moreau, M. Jimenez-Sanchez, S.Y. Breusegem, M.E. Harbour, and D.C. Rubinsztein. 2014. Mutation in VPS35 associated with Parkinson’s disease impairs WASH complex association and inhibits autophagy. Nat Commun. 5:3828.

Zhang, P., Y. Wu, T.Y. Belenkaya, and X. Lin. 2011. SNX3 controls Wingless/Wnt secretion through regulating retromer-dependent recycling of Wntless. Cell Res. 21:1677–1690.

Zhou, B., Y. Wu, and X. Lin. 2011. Retromer regulates apical-basal polarity through recycling Crumbs. Dev Biol. 360:87–95.

