## Supplemental materials for "Opposing functions for retromer and Rab11 in extracellular vesicle cargo traffic at presynaptic terminals"

Walsh et al.

Supplementary Figure Legends
3 Supplementary Figures
Table 1: Fly strains
Table 2: Antibodies and Reagents
Table 3: Statistics by Dataset

### Supplementary Figure Legends

**Figure S1: Presynaptic APP and Appl localize to NMJ EVs. (A)** Confocal images of NMJs from animals expressing muscle-driven ($GAL4^{BG57}$) UAS-APP-EGFP, stained with α-HRP. Arrows indicate presynaptically-derived HRP-positive puncta that do not co-localize with muscle-derived APP. (**B**) Schematic of domain organization and mutants in *Drosophila Appl*. (**C**) MaxIPs showing EV localization of *Drosophila* Appl. Yellow arrows indicate puncta that are positive for both Appl and α-HRP. White arrows indicate puncta that are solely α-HRP-positive. Cpx (presynaptic marker) is absent from Appl and α-HRP puncta. (**D**) MaxIP showing EV localization of *Drosophila* $Appl^{SD}$, and absence of α-Appl C-term antibody immunostaining for $Appl^{SD\Delta C}$. Scale bars are 5 μm; note that image display settings for α-Appl are different between (C) and (D) in order to visualize $Appl^{SD}$, which expresses at high levels. **Associated with Figure 1.**

**Figure S2: Neuronal retromer co-localizes with APP and restricts APP compartment size.** (**A**) APP-containing compartments are larger in *Vps35* mutants. MaxIPs and quantification of representative SIM images of NMJs expressing APP-EGFP in the indicated genotypes. Scale bar is 2 μm. (**B**) The fraction of postsynaptic APP-EGFP versus total APP-EGFP at *Vps35* mutant NMJs is not strongly correlated. (**C**) The fraction of APP-EGFP that is postsynaptic slightly increases in *Vps35* NMJs. (**D**) Vps35 exhibits both pre- and post-synaptic punctate localization and is enriched in perinuclear puncta in the muscle. MaxIPs of NMJ or muscle nucleus (bottom right) expressing Vps35-TagRFPt from the endogenous *Vps35* locus. Scale bar is 5 μm for NMJ and 10 μm for muscle. (**E**) Presynaptic APP-EGFP and Vps35-HA exhibit partial co-localization, indicated by arrows. Shown are MaxIPs of structured illumination microscopy (SIM) images; scale bar is 2 μm. (**F**) Controls for Vps35-HA rescue**.** Neuronal UAS-Vps35-HA expression does not affect levels of APP-EGFP in a wild-type background. Measurements were normalized to presynaptic mean of control (green line). Bar graphs show mean +/- s.e.m.; dots show all data points representing individual NMJs. **See Table 3 for detailed genotypes and statistical tests. Associated with Figure 2**

**Figure S6. Lysosome dysfunction does not cause retromer-dependent EV cargo sorting defects. (A-D)** Retromer loss does not causes accumulation of lysosome-like compartments. (**A)** MaxIPs of NMJs from larvae expressing neuronally-driven Spin-GFP or ATG8-GFP. Control is $GAL4^{C155}$. (**B**) Quantification of (**A**), normalized to mean intensity of wild-type control (green line). (**C**) MaxIPs of NMJs and the muscle perinuclear region labeled with LysoTracker Deep Red. Muscles exhibit a decrease in LysoTracker intensity indicating a reduction in rather than an increase in acidification of the lysosomal compartment. Control is $w^{1118}$. (**D, E)** Mutants that disrupt lysosome function do not exhibit EV cargo accumulation or increased postsynaptic EV cargo. (**D**) MaxIPs of NMJs from larvae expressing endogenously-tagged Syt4-EGFP. Controls are $w^{1118}$. (**E**) Quantification of (**D**). (**F-G**) Pharmacological inhibition of lysosome function does not affect EV cargo levels. (**F**) MaxIPs of NMJs from larvae expressing endogenously-tagged Syt4-EGFP and treated with the indicated concentration of chloroquine (CQ). (**G**) Quantification of (**F**). All conditions normalized to control. Red line indicates mean Syt4-EGFP levels in Vps35 mutants from **Fig 2**. Scale bars are 5 µm. Bar graphs show mean +/- s.e.m.; dots show all data points representing individual NMJs. **Associated with Figure 6.**

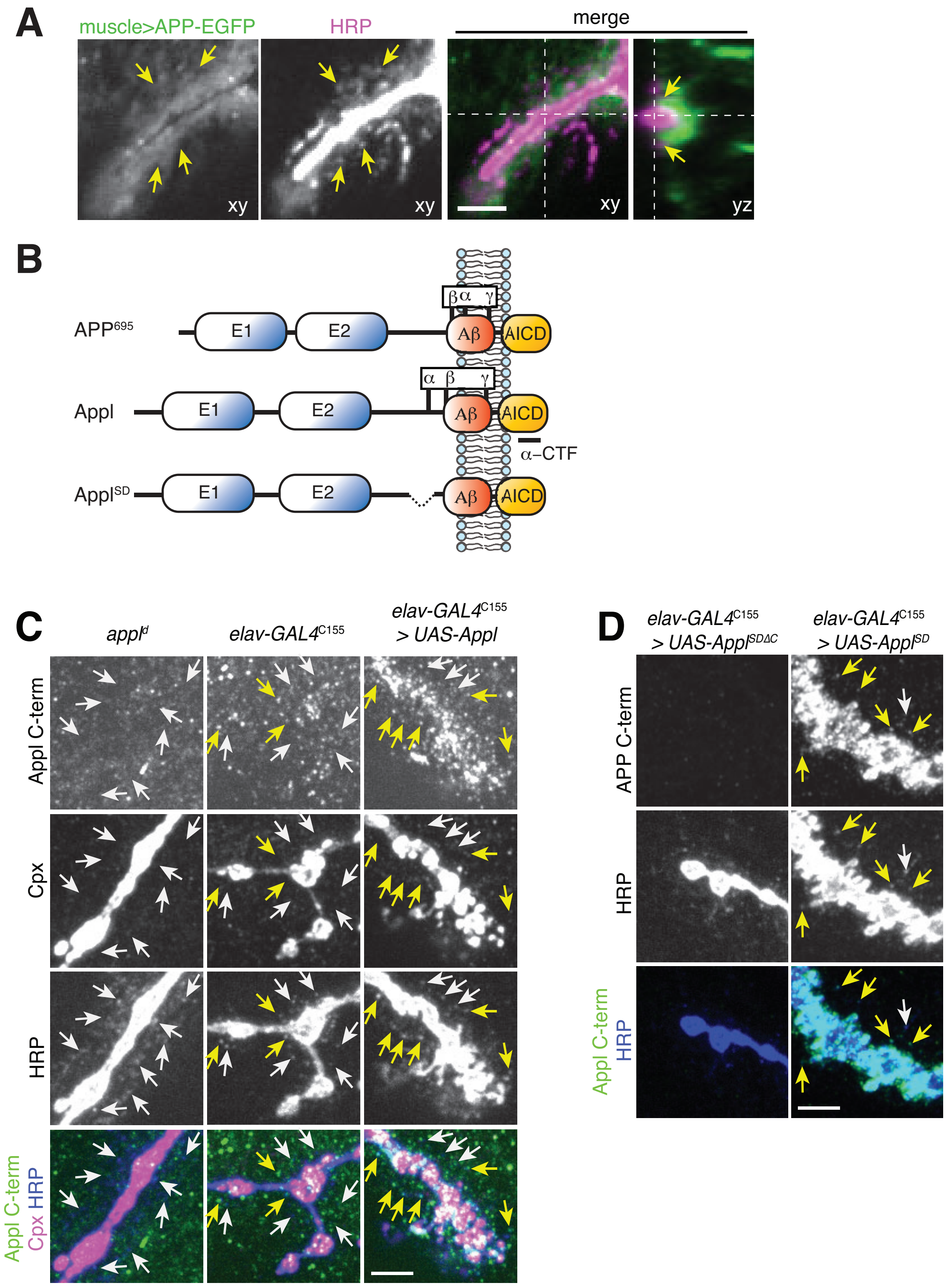
A
muscle>APP-EGFP
HRP
merge
xy
xy
xy
yz
B
APP695
E1
E2
Aβ
AICD
β α γ
Appl
α β γ
α–CTF
ApplSD
C
appld
elav-GAL4C155
elav-GAL4C155 > UAS-Appl
Appl C-term
Cpx
HRP
Appl C-term Cpx HRP
D
elav-GAL4C155 > UAS-ApplSDΔC
elav-GAL4C155 > UAS-ApplSD
APP C-term
HRP
Appl C-term HRP

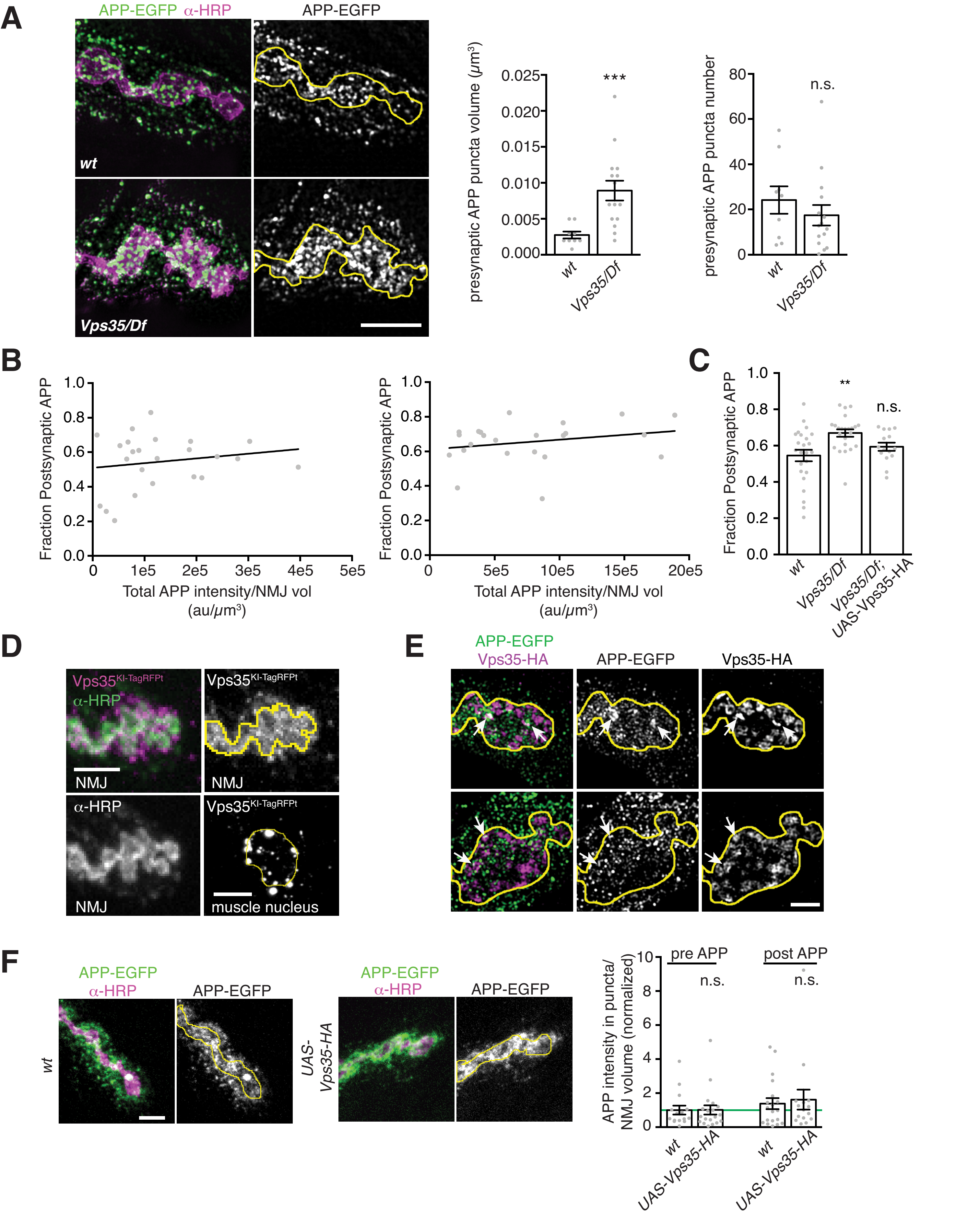
A
APP-EGFP α-HRP
APP-EGFP
wt
Vps35/Df
presynaptic APP puncta volume (μm³)
0.025
0.020
0.015
0.010
0.005
0.000
***
presynaptic APP puncta number
80
60
40
20
0
n.s.
B
Fraction Postsynaptic APP
1.0
0.8
0.6
0.4
0.2
0.0
0
1e5
2e5
3e5
4e5
5e5
Total APP intensity/NMJ vol
(au/μm³)
5e5
10e5
15e5
20e5
C
**
n.s.
Vps35/Df; UAS-Vps35-HA
D
Vps35KI-TagRFPt
α-HRP
NMJ
muscle nucleus
E
APP-EGFP
Vps35-HA
F
APP intensity in puncta/ NMJ volume (normalized)
10
8
6
4
2
pre APP
post APP
UAS-Vps35-HA

Walsh et al. Figure 6 Supplement

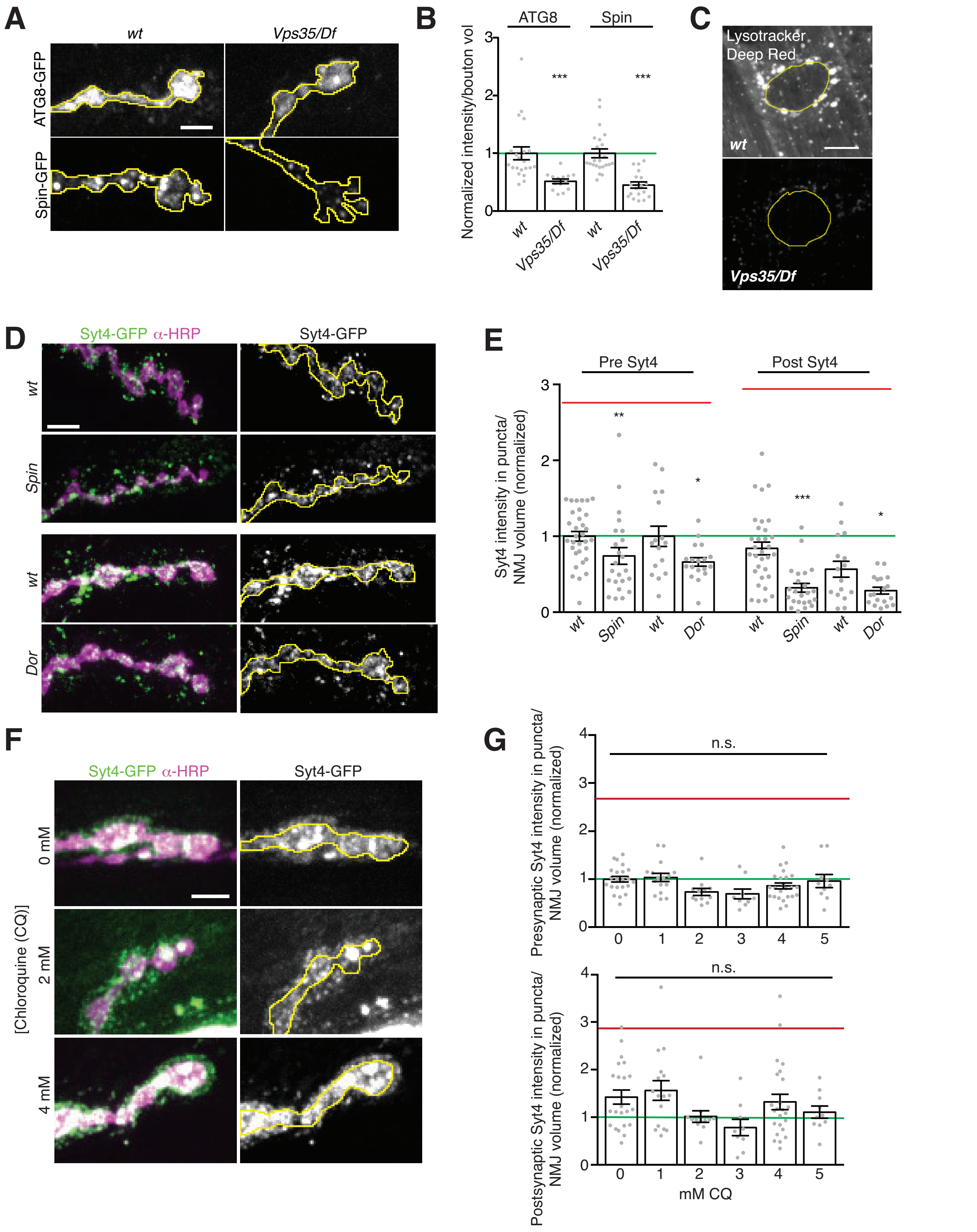

**TABLE 1: Fly Strains**

| Experimental Models: Organism/allele | | |
|---|---|---|
| *D. melanogaster:* $GAL4^{C155}$ | (Lin and Goodman, 1994) | Flybase ID: FBti0002575 BL548 |
| *D. melanogaster: w[1118]* | (Hazelrigg et al., 1984) | Flybase ID: FBal0018186 |
| *D. melanogaster:* UAS-Appl | (Torroja et al., 1999) | Flybase ID: FBti0128472 BL38403 |
| *D. melanogaster:* $UAS\text{-}Appl^{SD}$ | (Torroja et al., 1999) | Flybase ID: FBal0103093 BL29863 |
| *D. melanogaster*:{pUASt-Tsp43Ej-3xHA}AttP1 | Vivian Budnik | |
| *D. melanogaster:* $appl^{d}$ | (Luo et al., 1992) | Flybase ID: FBal0028990 BL43632 |
| *D. melanogaster:* $Vps35^{e42}$ | (Port et al., 2008) | Flybase ID: FBal0221801 |
| *D. melanogaster:* UAS-Tkv-mCherry | (Deshpande et al., 2016) | Flybase ID: FBal0322957 |
| *D. melanogaster*:{pUASt-Evi-3xHA}AttP2 | (Koles et al., 2012) | Flybase ID: FBal0269140 |
| *D. melanogaster:* Syt4-EGFP-KI | This work | |
| *D. melanogaster:* Df(2R)Exel6078 (removes *Vps35*) | (Parks et al., 2004) | Flybase ID: FBab0038016 BL7558 |
| *D. melanogaster:* $Vps26^{G2008}$ | (Linhart et al., 2014) | Flybase ID: FBal0220056 BL26623 |
| *D. melanogaster:* $Snx1^{\Delta 2}$ | (Zhang et al., 2011) | Flybase ID: FBal0336681 |
| *D. melanogaster:* Df(2L)Exel7016 (removes *Snx1*) | (Parks et al., 2004) | Flybase ID: FBab0037900 BL7787 |
| *D. melanogaster:* $Snx3^{\Delta 1}$ | (Zhang et al., 2011) | Flybase ID: FBal0267851 |
| *D. melanogaster:* Df(3R)exel7317 (removes *Snx3*) | (Parks et al., 2004) | Flybase ID: FBst0007932 BL7932 |

| | | |
|---|---|---|
| *D. melanogaster: Snx6*$^{1}$ | (Zhang et al., 2011) | Flybase ID: FBal 0267856 |
| *D. melanogaster:* Df(2L)BSC227 (removes *Snx6*) | | Flybase ID: FBab0045009 BL9704 |
| *D. melanogaster:* PBac{UAS-Vps35-HA} | (Wang et al., 2014) | Flybase ID: FBti181537 BL67152 |
| *D. melanogaster:* Vps35-TagRFPt$^{KI}$ | (Koles et al., 2015) | Flybase ID: FBal0321557 BL66527 |
| *D. melanogaster: Vps35*-TagRFPt$^{D628N\text{-}KI}$ | (Koles et al., 2015) | Flybase ID: FBal0321559 BL66528 |
| *D. melanogaster:* GAL4$^{C380}$ | (Budnik et al., 1996) | Flybase ID: FBti0016294 |
| *D. melanogaster:* GAL4$^{C57}$ | (Budnik et al., 1996) | Flybase ID: FBti0016293 |
| *D. melanogaster:* GFP-Rab5-KI | (Fabrowski et al., 2013) | Flybase ID: FBal0288693 |
| *D. melanogaster:* YFP-Myc-Rab7KI | (Dunst et al., 2015) | Flybase ID: FBal0314198 |
| *D. melanogaster:* UAS-Rab5$^{S43N}$ (II) | (Entchev et al., 2000) | Flybase ID: FBal0120656 BL42703 |
| *D. melanogaster:* UAS-Rab5$^{Q88L}$(III) | (Shimizu et al., 2003) | Flybase ID: FBal0182043 BL43335 |
| *D. melanogaster:* UAS-Rab7$^{Q67L}$(III) | (Zhang et al., 2007) | Flybase ID FBal0215400 BL9779 |
| *D. melanogaster:* UAS-Rab7$^{T22N}$ (III) | (Assaker et al., 2010) | Flybase ID: FBal0248958 BL9778 |
| *D. melanogaster:* UAS-Rab11$^{N124I}$ (II) | (Satoh et al., 2005) | Flybase ID FBal0190955 |
| *D. melanogaster:* UAS-myc-Spin-GFP | (Sweeney and Davis, 2002) | FlyBase ID FBal0143205 BL39668 |
| *D. melanogaster:* UAS-ATG8a-GFP (II) | (Juhasz et al., 2008) | FlyBase ID FBal0221640 |

| | | |
|---|---|---|
| | | BL52005 |
| *D. melanogaster: Spin*$^5$ | (Sweeney and Davis, 2002) | FlyBase ID FBal0064248 |
| *D. melanogaster: Spin*$^4$ | (Sweeney and Davis, 2002) | FlyBase ID FBal0008147 |
| *D. melanogaster: dor*$^4$ | (Belyaeva et al., 1980) | FlyBase ID FBal0002732 BL35 |
| *D.melanogaster: rab11*$^{ex2}$ | (Dollar et al., 2002) | Flybase ID FBal0135840 |
| *D. melanogaster: rab11*$^{93Bi}$ | (Eisenberg et al., 1990) | Flybase ID FBal0031959 |

Bloomington Drosophila Stock Center stock numbers denoted with BL.

**References**


Assaker, G., D. Ramel, S.K. Wculek, M. Gonzalez-Gaitan, and G. Emery. 2010. Spatial restriction of receptor tyrosine kinase activity through a polarized endocytic cycle controls border cell migration. *Proc Natl Acad Sci U S A*. 107:22558-22563.

Belyaeva, E.S., M.G. Aizenzon, V.F. Semeshin, Kiss, II, K. Koczka, E.M. Baritcheva, T.D. Gorelova, and I.F. Zhimulev. 1980. Cytogenetic analysis of the 2B3-4--2B11 region of the X-chromosome of *Drosophila* melanogaster. I. Cytology of the region and mutant complementation groups. *Chromosoma*. 81:281-306.

Budnik, V., Y.H. Koh, B. Guan, B. Hartmann, C. Hough, D. Woods, and M. Gorczyca. 1996. Regulation of synapse structure and function by the *Drosophila* tumor suppressor gene dlg. *Neuron*. 17:627-640.

Deshpande, M., Z. Feiger, A.K. Shilton, C.C. Luo, E. Silverman, and A.A. Rodal. 2016. Role of BMP receptor traffic in synaptic growth defects in an ALS model. *Mol Biol Cell*. 27:2898-2910.

Dollar, G., E. Struckhoff, J. Michaud, and R.S. Cohen. 2002. Rab11 polarization of the *Drosophila* oocyte: a novel link between membrane trafficking, microtubule organization, and oskar mRNA localization and translation. *Development*. 129:517-526.

Dunst, S., T. Kazimiers, F. von Zadow, H. Jambor, A. Sagner, B. Brankatschk, A. Mahmoud, S. Spannl, P. Tomancak, S. Eaton, and M. Brankatschk. 2015. Endogenously tagged rab proteins: a resource to study membrane trafficking in *Drosophila*. *Dev Cell*. 33:351-365.

Eisenberg, M., K. Gathy, T. Vincent, and J. Rawls. 1990. Molecular cloning of the UMP synthase gene rudimentary-like from *Drosophila melanogaster*. *Mol Gen Genet*. 222:1-8.

Entchev, E.V., A. Schwabedissen, and M. Gonzalez-Gaitan. 2000. Gradient formation of the TGF-beta homolog Dpp. *Cell*. 103:981-991.

Fabrowski, P., A.S. Necakov, S. Mumbauer, E. Loeser, A. Reversi, S. Streichan, J.A. Briggs, and S. De Renzis. 2013. Tubular endocytosis drives remodelling of the apical surface during epithelial morphogenesis in *Drosophila*. *Nat Commun*. 4:2244.

Hazelrigg, T., R. Levis, and G.M. Rubin. 1984. Transformation of white locus DNA in *Drosophila*: dosage compensation, zeste interaction, and position effects. *Cell*. 36:469-481.

Juhasz, G., J.H. Hill, Y. Yan, M. Sass, E.H. Baehrecke, J.M. Backer, and T.P. Neufeld. 2008. The class III PI(3)K Vps34 promotes autophagy and endocytosis but not TOR signaling in *Drosophila*. *J Cell Biol*. 181:655-666.

Koles, K., E.M. Messelaar, Z. Feiger, C.J. Yu, C.A. Frank, and A.A. Rodal. 2015. The EHD protein Past1 controls postsynaptic membrane elaboration and synaptic function. *Mol Biol Cell*. 26:3275-3288.

Koles, K., J. Nunnari, C. Korkut, R. Barria, C. Brewer, Y. Li, J. Leszyk, B. Zhang, and V. Budnik. 2012. Mechanism of evenness interrupted (Evi)-exosome release at synaptic boutons. *J Biol Chem*. 287:16820-16834.

Lin, D.M., and C.S. Goodman. 1994. Ectopic and increased expression of Fasciclin II alters motoneuron growth cone guidance. *Neuron*. 13:507-523.

Linhart, R., S.A. Wong, J. Cao, M. Tran, A. Huynh, C. Ardrey, J.M. Park, C. Hsu, S. Taha, R. Peterson, S. Shea, J. Kurian, and K. Venderova. 2014. Vacuolar protein sorting 35 (Vps35) rescues locomotor deficits and shortened lifespan in *Drosophila* expressing a Parkinson's disease mutant of Leucine-Rich Repeat Kinase 2 (LRRK2). *Mol Neurodegener*. 9:23.

Luo, L., T. Tully, and K. White. 1992. Human amyloid precursor protein ameliorates behavioral deficit of flies deleted for Appl gene. *Neuron*. 9:595-605.

Parks, A.L., K.R. Cook, M. Belvin, N.A. Dompe, R. Fawcett, K. Huppert, L.R. Tan, C.G. Winter, K.P. Bogart, J.E. Deal, M.E. Deal-Herr, D. Grant, M. Marcinko, W.Y. Miyazaki, S. Robertson, K.J. Shaw, M. Tabios, V. Vysotskaia, L. Zhao, R.S. Andrade, K.A. Edgar, E. Howie, K. Killpack, B. Milash, A. Norton, D. Thao, K. Whittaker, M.A. Winner, L. Friedman, J. Margolis, M.A. Singer, C. Kopczynski, D. Curtis, T.C. Kaufman, G.D. Plowman, G. Duyk, and H.L. Francis-Lang. 2004. Systematic generation of high-resolution deletion coverage of the *Drosophila melanogaster* genome. *Nat Genet*. 36:288-292.

Port, F., M. Kuster, P. Herr, E. Furger, C. Banziger, G. Hausmann, and K. Basler. 2008. Wingless secretion promotes and requires retromer-dependent cycling of Wntless. *Nat Cell Biol*. 10:178-185.

Satoh, A.K., J.E. O'Tousa, K. Ozaki, and D.F. Ready. 2005. Rab11 mediates post-Golgi trafficking of rhodopsin to the photosensitive apical membrane of *Drosophila* photoreceptors. *Development*. 132:1487-1497.

Shimizu, H., S. Kawamura, and K. Ozaki. 2003. An essential role of Rab5 in uniformity of synaptic vesicle size. *J Cell Sci*. 116:3583-3590.

Sweeney, S.T., and G.W. Davis. 2002. Unrestricted synaptic growth in spinster-a late endosomal protein implicated in TGF-beta-mediated synaptic growth regulation. *Neuron*. 36:403-416.

Torroja, L., H. Chu, I. Kotovsky, and K. White. 1999. Neuronal overexpression of APPL, the *Drosophila* homologue of the amyloid precursor protein (APP), disrupts axonal transport. *Curr Biol*. 9:489-492.

Wang, S., K.L. Tan, M.A. Agosto, B. Xiong, S. Yamamoto, H. Sandoval, M. Jaiswal, V. Bayat, K. Zhang, W.L. Charng, G. David, L. Duraine, K. Venkatachalam, T.G. Wensel, and H.J. Bellen. 2014. The retromer complex is required for rhodopsin recycling and its loss leads to photoreceptor degeneration. *PLoS Biol*. 12:e1001847.

Zhang, J., K.L. Schulze, P.R. Hiesinger, K. Suyama, S. Wang, M. Fish, M. Acar, R.A. Hoskins, H.J. Bellen, and M.P. Scott. 2007. Thirty-one flavors of *Drosophila* rab proteins. *Genetics*. 176:1307-1322.

Zhang, P., Y. Wu, T.Y. Belenkaya, and X. Lin. 2011. SNX3 controls Wingless/Wnt secretion through regulating retromer-dependent recycling of Wntless. *Cell Res*. 21:1677-1690.

**TABLE 2: ANTIBODIES**

| REAGENT | SOURCE | IDENTIFIER | Concentration |
|---|---|---|---|
| **Antibodies** | | | |
| α-HRP-488, 549, 647 | Jackson Immunoresearch | | 1:250-1:500 (IHC) |
| α–GFP nanobody | Nanotag Biotechnologies | N0304 | 1:250 (IHC) |
| α-HA | Covance MMS-101R | HA.11/16B12 | 1:500 (IHC) |
| α-Cpx | (Huntwork and Littleton, 2007) | | 1:1000 (IHC) |
| α-Aβ (17-24) | Covance SIG-39220 | 4G8 | 1:50(IHC) |
| α-GFP | MBL International Corporation | MBL598 | 1:1000 (IB) |
| α-Appl-C-term | (Swanson et al., 2005) | | 1:1000 (IHC) |
| α-Rab11 | BD Biosciences | 610657 | 1:100 (IHC) |
| α-Synaptotagmin-1 | (West et al., 2015) | | 1:1000 (IHC) |

***References***

Huntwork, S., and J.T. Littleton. 2007. A complexin fusion clamp regulates spontaneous neurotransmitter release and synaptic growth. *Nat Neurosci*. 10:1235-1237.

Swanson, T.L., L.M. Knittel, T.M. Coate, S.M. Farley, M.A. Snyder, and P.F. Copenhaver. 2005. The insect homologue of the amyloid precursor protein interacts with the heterotrimeric G protein Go alpha in an identified population of migratory neurons. *Dev Biol*. 288:160-178.

West, R.J., Y. Lu, B. Marie, F.B. Gao, and S.T. Sweeney. 2015. Rab8, POSH, and TAK1 regulate synaptic growth in a *Drosophila* model of frontotemporal dementia. *J Cell Biol*. 208:931-947.

**TABLE 3: Statistics by dataset**

**Measurements key:**
Statistical tests were conducted relative to controls that were dissected and imaged in parallel. Matched controls and experiments are shown in the same table cell.
Presynaptic volume: α–HRP objects > 7 μm$^3$
Postsynaptic volume: 3 μm dilated from presynaptic volume
a: Sum intensity of signal in thresholded objects in presynaptic volume, normalized to presynaptic volume.
b: Sum intensity of signal in thresholded objects in postsynaptic volume, normalized to presynaptic volume.
SIM: structured illumination microscopy
SDCM: spinning disk confocal microscopy
EM: electron microscopy

| **Panel** | **Genotypes** | **n** | **Measurements (channel)** | **Statistical Test** |
|---|---|---|---|---|
| 2B (SDCM) | GAL4$^{C155}$/Y; UAS-APP-EGFP/+<br><br>GAL4$^{C155}$; UAS-APP-EGFP,*Vps35*$^{e42}$/ Df(Exel)6078<br><br>GAL4$^{C155}$/Y; UAS-APP-EGFP,*Vps35*$^{e42}$/ Df(Exel)6078; UAS-Vps35-HA/+ | 24 NMJs from 12 animals<br><br>23 NMJs from 16 animals<br><br>15 NMJs from 5 animals | a, b (APP-EGFP) | Presynaptic: Kruskal-Wallis ($p < 0.0001$)<br><br>Postsynaptic: Kruskal-Wallis ($p < 0.0001$) |
| 2C (SDCM) | GAL4$^{C155}$/Y; UAS-APP-EGFP/+<br><br>GAL4$^{C155}$; UAS-APP-EGFP,*Vps35*$^{e42}$/ Df(Exel)6078<br><br>GAL4$^{C155}$/Y; UAS-APP-EGFP,*Vps35*$^{e42}$/ Df(Exel)6078; UAS-Vps35-HA/+ | Same dataset as 2B | Overall mean in presynaptic volume (APP-EGFP; no APP threshold) | Kruskal-Wallis ($p<0.0001$) |
| 2E (SDCM) | GAL4$^{C155}$/Y; UAS-APP-EGFP/+<br><br>GAL4$^{C155}$; UAS-APP-EGFP,*Vps35*$^{e42}$/ Df(Exel)6078 | 30 axons, 56 cell bodies from 9 animals<br><br>33 axons, 56 cell bodies from 8 animals | axon measurements of sum intensity projection in axon only ( APP-EGFP)<br><br>cell body measurements from single slice through the middle of each cell body (APP-EGFP) | Axon: Mann-Whitney ($p < 0.0001$)<br><br>Cell bodies: Mann-Whitney ($p < 0.0001$) |
| 2G-I (SDCM) | $w^{1118}$/Y; ; Syt4-EGFP/+ (control for Vps35)<br><br>$w^{1118}$/Y; *Vps35*$^{e42}$/Df(Exel)6078; Syt4-EGFP/+ | 22 NMJs from 6 animals<br><br>17 NMJS from 6 animals | a, b (Syt4-EGFP) | Presynaptic: t-Test ($p < 0.0001$)<br><br>Postsynaptic: t-Test ($p < 0.0001$) |
| | $w^{1118}$/Y; ; Syt4-EGFP/+ (control for Vps26) | 15 NMJS from 3 animals | a, b (Syt4-EGFP) | Presynaptic: t-Test |

| | | | | |
|---|---|---|---|---|
| | *Vps26*$^{G2008}$/Y; ; Syt4-EGFP/+ | 14 NMJs from 4 animals | | (p = 0.0019)<br><br>Postsynaptic: t-Test (p < 0.0001) |
| | GAL4$^{C155}$/Y; UAS-Tkv-mCh/+;<br><br>GAL4$^{C155}$/Y; *Vps35*$^{e42}$, UAS-Tkv-mCh/ Df(Exel)6078; | 18 NMJs from 7 animals<br><br>15 NMJs from 6 animals | a, b (Tkv-mCh) | Presynaptic: Mann-Whitney (p = 0.0075)<br><br>Postsynaptic: Mann-Whitney (p < 0.0001) |
| | *w*$^{1118}$/Y<br><br>*w*$^{1118}$/Y; *Vps35*$^{e42}$/ Df(Exel)6078 | 22 NMJs from 8 animals<br><br>26 NMJs from 8 animals | a (Syt1) | Presynaptic: Mann-Whitney (p=0.2354) |
| Fig S2D (SDCM) | GAL4$^{C155}$/Y; UAS-APP-EGFP/+;<br><br>GAL4$^{C155}$/Y; UAS-APP-EGFP/+;UAS-Vps35-HA/+ | 15 NMJs from 5 animals<br><br>19 NMJs from 6 animals | a, b (APP-EGFP) | Presynaptic: Mann-Whitney (p = 0.8842)<br><br>Postsynaptic: Mann-Whitney (p = 0.7010) |
| Fig, S2F (SIM) | GAL4$^{C155}$/Y; UAS-APP-EGFP/+<br><br>GAL4$^{C155}$; UAS-APP-EGFP,*Vps35*$^{e42}$/ Df(Exel)6078 | 9 NMJs from 4 animals<br><br>15 NMJs from 6 animals | Volume of APP-thresholded objects within presynaptic volume | t-test (p = 0.0108) |
| Fig S2G (SIM) | GAL4$^{C155}$/Y; UAS-APP-EGFP/+<br><br>GAL4$^{C155}$; UAS-APP-EGFP,*Vps35*$^{e42}$/ Df(Exel)6078 | Same dataset as S2F | Number of APP-thresholded objects within presynaptic volume | t-test (p = 0.0715) |
| Fig S2H (SDCM) | GAL4$^{C155}$/Y; UAS-APP-EGFP/+; | Same dataset as Fig 2B | post APP-EGFP (b) vs. fraction APP extraneuronal | Linear regression, F test if slope>0 $R^2$ = 0.02990 p = 0.4191 |
| Fig S2I (SDCM) | GAL4$^{C155}$/Y; UAS-APP-EGFP,*Vps35*$^{e42}$/ Df(Exel)6078/ | Same dataset as Fig 2B | post APP-EGFP (b) vs. fraction APP extraneuronal | Linear regression, F test if slope>0 $R^2$ = 0.06021 p = 0.2591 |
| Fig S2J (SDCM) | GAL4$^{C155}$/Y; UAS-APP-EGFP/+<br><br>GAL4$^{C155}$; UAS-APP-EGFP,*Vps35*$^{e42}$/ Df(Exel)6078<br><br>GAL4$^{C155}$/Y; UAS-APP- | Same dataset as Fig 2B | APP in postsynaptic region/Total APP | Kruskal-Wallis (p<0.0024) |

| | | | | |
|---|---|---|---|---|
| | EGFP,*Vps35*$^{e42}$/ Df(Exel)6078;<br>UAS-Vps35-HA/+ | | | |
| Fig 3B<br>(EM) | $w^{1118}$/Y<br><br>$w^{1118}$/Y;<br>*Vps35*$^{e42}$/ Df(Exel)6078 | 18 boutons from 6 animals<br><br>28 boutons from 6 animals | Within 1 µm region of the bouton | Mann-Whitney<br>(p < 0.0001) |
| Fig 3C<br>(EM) | $w^{1118}$/Y<br><br>$w^{1118}$/Y;<br>*Vps35*$^{e42}$/ Df(Exel)6078 | Same dataset as Fig 3B | Inside the bouton | Mann-Whitney<br>(p = 0.3242) |
| Fig 3D<br>(EM) | $w^{1118}$/Y<br><br>$w^{1118}$/Y;<br>*Vps35*$^{e42}$/ Df(Exel)6078 | Same dataset as Fig 3B | Inside the bouton | Mann-Whitney<br>(p = 06891) |
| Fig 3E<br>(EM) | $w^{1118}$/Y<br><br>$w^{1118}$/Y;<br>*Vps35*$^{e42}$/ Df(Exel)6078 | Same dataset as Fig 3B | Inside the bouton | Mann-Whitney<br>(p = 0.3884) |
| Fig 4A<br>(SDCM) | $w^{1118}$/Y; ;<br><br>$w^{1118}$/Y; *Vps35*$^{e42}$/ Df(Exel)6078<br><br>$w^{1118}$; *Vps35*$^{WT\text{-}KI}$<br><br>$w^{1118}$; *Vps35*$^{D628N\text{-}KI}$<br><br>$w^{1118}$/Y; ; *Snx3*$^{\Delta 1}$/Df(Exel)7317<br><br>$w^{1118}$/Y; *Snx6*$^{\Delta 1}$, *Snx1*$^{\Delta 2}$/<br>Df(2L)BSC227, Df(Exel)7016<br><br>*Snx27*$^{KO}$/Y | 20 NMJs from 10 animals<br><br>21 NMJs from 15 animals<br><br>12 NMJs from 8 animals<br><br>15 NMJs from 8 animals<br><br>19 NMJs from 11 animals<br><br>21 NMJs from 9 animals<br><br>11 NMJs from 4 animals | Type 1b bouton number<br>Muscle 4,<br>Segment A3 | ANOVA<br>(p < 0.0001) |
| Fig 5A<br>(SDCM) | GAL4$^{C155}$/Y; UAS-APP-EGFP,*Vps35*$^{e42}$/*Vps35*$^{KI}$<br><br>GAL4$^{C155}$/Y; UAS-APP-EGFP,*Vps35*$^{e42}$/*Vps35*$^{D628N\text{-}KI}$ | 16 NMJs from 4 animals<br><br>16 NMJs from 4 animals | a, b (APP-EGFP) | Presynaptic:<br>Mann-Whitney<br>(p = 0.0075)<br><br>Postsynaptic:<br>Mann-Whitney<br>(p = 0.0426) |
| Fig 5B<br>(SDCM) | GAL4$^{C155}$/Y; UAS-APP-EGFP/+;<br><br>*Snx27*$^{KO}$, GAL4$^{C155}$/Y; UAS-APPEGFP/+ | 28 NMJs from 9 animals<br><br>22 NMJS from 8 animals | a, b (APP-EGFP) | Presynaptic:<br>Mann-Whitney<br>(p =0.8389)<br><br>Postsynaptic:<br>Mann-Whitney<br>(p = 0.6489) |

| | | | | |
|---|---|---|---|---|
| Fig 5C (SDCM) | GAL4[C155]/Y; UAS-APP-EGFP/+;<br><br>GAL4[C155]/Y; *Snx6*[1], UAS-APPEGFP, *Snx1*[Δ2]/ Df(2L)BSC227, Df(Exel)7016 | 24 NMJs from 10 animals<br><br>25 NMJS from 9 animals | a, b (APP-EGFP) | Presynaptic: Mann-Whitney ($p = 0.0032$)<br><br>Postsynaptic: T-test ($p = 0.0019$) |
| Fig 6A-B (SDCM) | *w*[1118]/Y;<br><br>*w*[1118]/Y; *Vps35*[e42]/Df(Exel)6078 | 13 NMJs from6 animals<br><br>19 NMJs from6 animals | a (α-Rab11) | T-Test ($p = 0.1550$) |
| | *w*[1118]/Y; YFP-myc-Rab5-KI/+<br><br>*w*[1118]/Y; *Vps35*[e42],YFP-myc-Rab5-KI/ Df(Exel)6078 | 18 NMJs from 5 animals<br><br>16 NMJS from 6 animals | a (GFP-Rab5) | Mann-Whitney ($p = 0.0001$) |
| | *w*[1118]/Y;YFP-myc-Rab7-KI/+<br><br>*w*[1118]/Y; *Vps35*[e42]/Df(Exel)6078; YFP-myc-Rab7-KI/+ | 17 NMJs from 6 animals<br><br>13 NMJs from 4 animals | a (YFP-Rab7) | Mann-Whitney ($p < 0.0001$) |
| Fig 6C-D (SDCM) | All datasets same as 6A-B | All datasets same as 6A-B | Rab-thresholded objects within presynaptic volume | Rab5 Density: T-test ($p < 0.0001$)<br><br>Rab5 Size Mann-Whitney ($p = 0.2572$)<br><br>Rab7 Density: T-test ($p = 0.0292$)<br><br>Rab7 Size: T-test ($p = 0.0024$) |
| Fig 6E-G (SIM) | GAL4[C155]/Y; UAS-APP-EGFP/+<br><br>GAL4[C155]/Y; Df(Exel)6078/UAS-APP-EGFP,*Vps35*[e42] | 9 NMJs from 5 animals<br><br>15 NMJs from 6 animals | α–Rab11-thresholded objects within presynaptic volume | Puncta Density: T-test ($p = 0.0715$)<br><br>Puncta Size T-test ($p = 0.0108$) |
| Fig S6A-B (SDCM) | GAL4[C155]/Y; UAS-ATG8-EGFP/+<br><br>GAL4[C155]/Y; Df(Exel)6078/*Vps35*[e42,]UAS-ATG8-EGFP | 21 NMJs from 6 animals<br><br>14 NMJs from 4 animals | a (Atg8-GFP) | Mann-Whitney ($p < 0.0001$) |

| | | | | |
|---|---|---|---|---|
| | GAL4$^{C155}$/Y; UAS-Spin-myc-EGFP/+<br><br>GAL4$^{C155}$/Y; Df(Exel)6078/*Vps35*$^{e42}$; UAS-Spin-myc-EGFP/+ | 23 NMJs from 7 animals<br><br>18 NMJs from 7 animals | a (Spin-GFP) | T-test<br>($p = 0.002$) |
| Fig S6D-E (SDCM) | *w*$^{1118}$/Y; ; Syt4-EGFP/+<br><br>*w*$^{1118}$/Y;*Spin*$^{4}$/*Spin*$^{5}$; Syt4-EGFP/+ | 32 NMJs from 11 animals<br><br>23 NMJs from 7 animals | a, b (Syt4-EGFP) | Presynaptic: Mann-Whitney ($p = 0.0058$)<br><br>Postsynaptic: Mann-Whitney ($p < 0.001$) |
| | *w*$^{1118}$/Y; Syt4-EGFP/+<br><br>*dor*$^{4}$/Y; ; Syt4-EGFP/+ | 16 NMJs from 6 animals<br><br>18 NMJs from 6 animals | a, b (Syt4-EGFP) | Presynaptic: T-test ($p = 0.0214$)<br><br>Postsynaptic: T-test ($p = 0.0154$) |
| Fig S6 F-G (SDCM) | (All genotypes *w*$^{1118}$/Y;Syt4-EGFP/+ or *w*$^{1118}$, Syt4-EGFP/+))<br>Control (0 mM)<br><br>1 mM<br><br>2 mM<br><br>3 mM<br><br>4 mM<br><br>5 mM | <br><br>22 NMJs from6 animals<br><br>16 NMJs from7 animals<br><br>12 NMJs from5 animals<br><br>9 NMJs from3 animals<br><br>24 NMJs from 7 animals<br><br>10 NMJs from3 animals | a, b (Syt4-EGFP) | Presynaptic: Kruskal-Wallis ($p = 0.0069$)<br><br>Postsynaptic: Kruskal-Wallis ($p = 0.0681$) |
| Fig 7A-C (SDCM) | GAL4$^{C380}$/Y; ; Syt4-EGFP/+<br><br>GAL4$^{C380}$/Y; UAS-Rab5$^{CA/+}$; Syt4-EGFP/+<br><br>GAL4$^{C380}$/Y; UAS-Rab5$^{DN}$/+; Syt4-EGFP/+<br><br>GAL4$^{C380}$/Y; UAS-Rab7$^{CA}$/+; Syt4-EGFP/+<br><br>GAL4$^{C380}$/Y; UAS-Rab7$^{DN}$/+; Syt4-EGFP/+ | 18 NMJs from 6 animals<br><br>12 NMJs from 4 animals<br><br>16 NMJs from 6 animals<br><br>22 NMJs from 7 animals<br><br>18 NMJs from 6 animals | a, b (Syt4-EGFP) | Presynaptic (7A): Kruskal-Wallis ($p < 0.0001$)<br><br>Postsynaptic (7A): Kruskal-Wallis ($p < 0.0001$) |
| | GAL4$^{C380}$/Y; ; Syt4-EGFP/+<br>Control for:<br><br>GAL4$^{C380}$/Y; UAS-Rab11$^{DN}$/+; Syt4-EGFP/+ | 22 NMJs from 7 animals<br><br>24 NMJs from 7 animals | a, b (Syt4-EGFP) | Presynaptic: Mann-Whitney ($p = 0.0021$)<br><br>Postsynaptic: |

| | | | | |
|---|---|---|---|---|
| | | | | Mann-Whitney ($p < 0.0001$) |
| | GAL4[C380]/Y; ; Syt4-EGFP/+ (7D) Control for:<br><br>GAL4[C380]/Y; UAS-Rab11[CA]/+; Syt4-EGFP/+ | 17 NMJs from 6 animals<br><br>18 NMJs from 6 animals | a, b (Syt4-EGFP) | Presynaptic: T-test ($p < 0.0001$)<br><br>Postsynaptic: T-test ($p < 0.0001$) |
| Fig 7D (SDCM) | GAL4[C155]/Y;UAS-APP-EGFP/+;<br><br>GAL4[C155]/Y;UAS-APP-EGFP/+; *Rab11*[ex2]/*Rab11*[93bi] | Same dataset as Fig 2B<br><br>23 NMJs from 10 animals | a, b (APP-EGFP) | Presynaptic: Mann-Whitney ($p < 0.0001$)<br><br>Postsynaptic Mann-Whitney ($p < 0.0001$) |
| Fig 7E (SDCM) | *w*[1118]/Y;YFP-myc-Rab7-KI/+<br><br>*w*[1118]/Y; *Vps35*[e42]/Df(Exel)6078; YFP-myc-Rab7-KI/+<br><br>*w*[1118]/Y; *Snx6*[Δ1], *Snx1*[Δ2]/Df(2L)BSC227, Df(Exel)7016; YFP-myc-Rab7-KI/+ | 24 NMJs from 7 animals<br><br>19 NMJs from 6 animals<br><br>29 NMJs from 7 animals | a (YFP-Rab7)<br><br>Puncta size and number: Rab-thresholded objects within presynaptic volume | ANOVA ($p < 0.0001$) |
| Fig 8A-B | GAL4[C155]/Y; UAS-APP-EGFP/+;<br><br>GAL4[C155]/Y; Df(Exel)6078/UAS-APP-EGFP,*Vps35*[e42]; | 21 NMJs from 10 animals<br><br>15 NMJs from 9 animals | Manders Coefficient | APP in Rab11 Mann-Whitney ($p = 0.1224$)<br><br>Rab11 in APP t-test ($p = 0.0107$) |
| Fig 8D-E | *w*[1118]/Y; ; Syt4-EGFP/+<br><br>*w*[1118]/Y; Df(Exel)6078/*Vps35*[e42]; Syt4-EGFP/+<br><br>*w*[1118]/Y; ; *Rab11*[ex2], Syt4-EGFP /*Rab11*[93Bi]<br><br>*w*[1118]/Y; Df(Exel)6078/*Vps35*[e42]; *Rab11*[ex2]; Syt4-EGFP /*Rab11*[93bi] | 16 NMJS from 8 animals<br><br>29 NMJs from 8 animals<br><br>18 NMJS from 9 animals<br><br>30 NMJs from 8 animals | a (Syt4-EGFP)<br><br>b (Syt4-EGFP) | Presynaptic: ANOVA ($p < 0.0001$)<br><br>Postsynaptic: Kruskal-Wallis ($p < 0.0001$) |
